# Metrics for Graph Comparison: A Practitioner’s Guide

**DOI:** 10.1101/611509

**Authors:** Peter Wills, François G. Meyer

## Abstract

Comparison of graph structure is a ubiquitous task in data analysis and machine learning, with diverse applications in fields such as neuroscience [1], cyber security [2], social network analysis [3], and bioinformatics [4], among others. Discovery and comparison of structures such as modular communities, rich clubs, hubs, and trees in data in these fields yields insight into the generative mechanisms and functional properties of the graph.

Often, two graphs are compared via a pairwise distance measure, with a small distance indicating structural similarity and vice versa. Common choices include spectral distances (also known as *λ* distances) and distances based on node affinities (such as `DeltaCon` [5]). However, there has of yet been no comparative study of the efficacy of these distance measures in discerning between common graph topologies and different structural scales.

In this work, we compare commonly used graph metrics and distance measures, and demonstrate their ability to discern between common topological features found in both random graph models and empirical datasets. We put forward a multi-scale picture of graph structure, in which the effect of global and local structure upon the distance measures is considered. We make recommendations on the applicability of different distance measures to empirical graph data problem based on this multi-scale view. Finally, we introduce the Python library `NetComp` which implements the graph distances used in this work.

## Introduction

In the era of big data, comparison and matching are ubiquitous tasks. A graph is a particular type of data structure which records the interactions between some collection of agents.^1^ This type of data structure relates connections between objects, rather than directly relating the properties of those objects. The interconnectedness of the object in graph data disallows many common statistical techniques used to analyze tabular datasets. The need for new analytical techniques for visualizing, comparing, and understanding graph data has given rise to a rich field of study [6].

In this work, we focus on tools for pairwise comparison of graphs. Such comparison often takes place within the contest of anomaly detection and graph matching. In the former, one has a sequence of graphs (often a time series) and hopes to establish at what time steps the graphs change “significantly” at any given time step. In the latter, one has a collection of graphs, and wants to establish whether a sample is likely to have been drawn from that collection. Both problems require the ability to effectively compare two graphs. However, the utility of any given comparison method varies with the type of information the user is looking for; one may care primarily about large scale graph features such as community structure or the existence of highly connected “hubs”; or, one may be focused on smaller scale structure such as local connectivity (i.e. the degree of a vertex) or the ubiquity of substructures such as triangles.

Existing surveys of graph distances are limited to observational datasets [7]. While authors try to choose datasets that are exemplars of certain classes of networks (e.g. social, biological, or computer networks), it is difficult to generalize these studies to other datasets.

In this paper, we take a different approach. We consider existing ensembles of random graphs as prototypical examples of certain graph *structures*, which are the building blocks of existing empirical network datasets. We propose therefore to study the ability of various distances to compare two samples randomly drawn from distinct ensembles of graphs. Our investigation is concerned with the relationship between the families of graph ensembles, the structural features characteristic of these ensembles, and the sensitivity of the distances to these characteristic structural features.

The myriad proposed techniques for graph comparison [8] are severely reduced in number when one requires the practical restriction that the algorithm run in a reasonable amount of time on large graphs. Graph data frequently consists of 10^4^ to 10^8^ vertices, and so algorithms whose complexity scales quadratically with the size of the graph quickly become unfeasible. In this work, we restrict our attention to approaches where the calculation time scaled linearly or near-linearly with the number of vertices in the graph for sparse graphs.^2^

In the past 40 years, many random graph models have been developed which emulate certain features found in real-world graphs [9, 10]. A rigorous probabilistic study of the application of graph distances to these random models is difficult because the models are often defined in terms of a generative process rather than a distribution over the space of possible graphs. As such, researchers often restrict their attention to very small, deterministic graphs (see e.g. [11]) or to very simple random models, such as that proposed by Erdős and Rényi [12]. Even in these simple cases, rigorous probabilistic analysis can be prohibitively difficult. We adopt a numerical approach, in which we sample from random graph distributions and observe the empirical performance of various distance measures.

Throughout the work, we understand the observed results through a lens of global versus local graph structure. Examples of global structure include community structure and the existence of well-connected vertices (often referred to as “hubs”). Examples of local structure include the median degree in the graph, or the density of substructures such as triangles. Our results demonstrate that some distances are particularly tuned towards observing global structure, while some naturally observe both scales. In both our empirical and numerical experiments, we use this multi-scale interpretation to understand why the distances perform the way they do on a given model, or on given empirical graph data.

The paper is structured as follows: in Section, we introduce the distances used, and establish the state of knowledge regarding each. In Section, we similarly introduce the random graph models of study and discuss their important features. In Section we numerically examine the ability of the distances to distinguish between the various random graph models. The reader who is already familiar with the graph models and distances discussed can skip to Section for a discussion of the results of our evaluation of the distances on the various random graph models, referencing the results in Section as necessary. In Section, we apply the distances to empirical graph data and discuss the results. Finally, Section summarizes the work and summarizes our recommendations. In the appendix, we introduce and discuss NetComp, the Python package which implements the distances used to compare the graphs throughout the paper.

### Graph Distance Measures

Let us begin by introducing the distances we will use in this study, and discussing the state of the knowledge for each. We have chosen both standard and cutting-edge distances, with the requirement that the algorithms be computable in a reasonable amount of time on large, sparse graphs. In practice, this means that the distances must scale linearly or near-linearly in the size in the graph.

We refer to these tools as “distance measures,” as many of them do not satisfy the technical requirements of a metric. Although all are symmetric, they may fail one or more of the other requirements of a mathematical metric. This can be very problematic if one hopes to perform rigorous analysis on these distances, but in practice it is generally not significant. Consider the requirement of identity of indiscernible, in which *d*(*G, G′*) = 0 if and only if *G* = *G′*. we rarely encounter two graphs where *d*(*G, G′*) = 0; we are more frequently concerned with an approximate form of this statement, in which we wish to deduce that *G* is similar to *G′* from the fact that *d*(*G, G′*) is small. Similarly, although the triangle inequality is foundational in approximation and proof methods in analysis, it is rarely employed in our process in applying these distances for anomaly detection.

### Notation

We must first introduce the notation used throughout the paper. It is standard wherever possible.

We denote by *G* = (*V, E, W*) a graph with vertex set *V* = *{*1, *…, n}* and edge set *E* ⊆ *V × V*. The function *W* : *E* → ***R***^+^ assigns each edge (*i, j*) in *E* a positive number, which we denote *w*_*i,j*_. We call *n* = |*V*| the **size** of the graph, and denote by 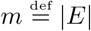 the number of edges. For *i ∈ V* and *j ∈ V*, we say *i* ~ *j* if (*i, j*) *∈ E*. The matrix ***A*** is called the **adjacency matrix**, and is defined as

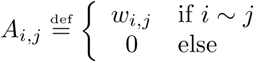

The **degree** *d*_*i*_ of a vertex is defined as 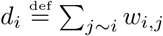. The **degree matrix *D*** is the diagonal matrix of degrees, so *D*_*i,i*_ = *d*_*i*_ and *D*_*i,j*_ = 0 for *i ≠ j*. The **Laplacian matrix** (or just Laplacian) of *G* is given by 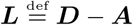. The **normalized Laplacian** is defined as 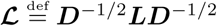, where the diagonal matrix ***D***^−1/2^ is given by

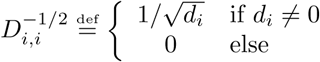

We refer to ***A***, ***L***, and **ℒ** as **matrix representations** of *G*. These are not the only useful matrix representation of a graph, although they are some of the most common. For a more diverse catalog of representations, see [13]. Note that other normalizations of the Laplacian matrix are possible; a popular choice is normalizing the rows so that they sum to one, which results in the transition matrix for a random walk on the graph. Our choice maintains symmetry, and thus strictly real eigenvalues and eigenvectors, a property which the row-normalized Laplacian lacks. A real spectrum simplifies computations, for example, when one wishes to form a basis of eigenvectors in order to decompose real-valued functions on the graph. However, the interpretability of the spectrum comes at the cost of interpretability of the matrix itself; while the row-stochastic normalization has an easy-to-understand function in terms of random walks, the interpretation of our normalized Laplacian **ℒ** is not so straightforward.

The **spectrum** of a matrix is the sorted sequence of eigenvalues. Whether the sequence is ascending or descending depends on the matrix in question. We denote the *i*^th^ eigenvalue of the adjacency matrix by 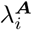, where 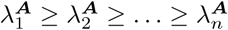. The *i*^th^ eigenvalue of the Laplacian matrix are denotes by 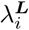, with the eigenvalues sorted in **ascending** order, so that 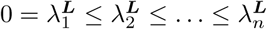. We similarly denote the *i*^th^ eigenvalue of the normalized Laplacian by 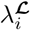, with 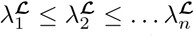.

Two graphs are **isomorphic** if and only if there exists a map between their vertex sets under which the two edge sets are equal. Since our vertex sets are integers, we can simplify this definition. In particular, let us say that *G ≅ G′* if and only if there exists a permutation matrix ***P*** such that ***A****′* = ***P*** ^*T*^ ***AP***.

We say that a distance *d* requires **node correspondence** when there exist graphs *G, G′*, and *H* such that *G ≅ G′* but *d*(*G, H*) ≠ *d*(*G′, H*). Intuitively, a distance requires node correspondence when one must know some meaningful mapping between the vertex sets of the graphs under comparison.

### Graph Distance Taxonomy

The distance functions we study divide naturally into two categories, which we will now describe. These categories are not exhaustive; many distance functions (including one we employ in our experiments) do not fit neatly into either category. Akoglu et al. [8] provide an alternative taxonomy; our taxonomy refines a particular group of methods they refer to as “feature-based”.^3^

#### Spectral Distances

Let us first discuss spectral distances, also known as *λ* distances. We will briefly review the necessary background; for a good introduction to spectral methods of graph comparison, see [13].

We will first define the adjacency spectral distance; the Laplacian and normalized Laplacian spectral distances are defines similarly. Let *G* and *G′* be graphs of size *n*, with adjacency spectra *λ*^***A***^ and 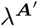, respectively. The **adjacency spectral distance** between the two graphs is defined as

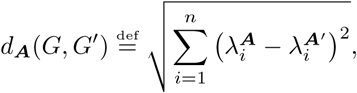

which is just the distance between the two spectra in the 𝓁_2_ metric. We could use any 𝓁_*p*_ metric here, for *p ∈* [0, *∞*]. The choice of *p* is informed by how much one wishes to emphasize outliers; in the limiting case of *p* = 0, the metric returns the measure of the set over which the two vectors are different, and when *p* = *∞* only the largest element-wise difference between the two vectors is returned. Note that for *p <* 1 the 𝓁_*p*_ distances are not true metrics (in particular, they fail the triangle inequality) but they still may provide valuable information. For a more detailed discussion on 𝓁_*p*_ norms, see [14].

The Laplacian and normalized Laplacian spectral distances *d*_***L***_ and *d*_***ℒ***_ are defined in the exact same way. In general, one can define a spectral distance for any matrix representation of a graph; for results on more than just the three we analyze here, see [13]. Spectral distances are invariant under permutations of the vertex labels; that is to say, if ***P*** is a permutation matrix, then the spectrum of ***A*** is equal to the spectrum of ***P*** ^*T*^ ***AP***. This allows us to directly compare the topological similarity of two graphs without having to discover any mapping between the vertex sets.

In practice, it is often the case that only the first *k* eigenvalues are compared, where *k* ≪ *n*. We refer to such truncated *λ* distances as *λ*_*k*_ distances. When using *λ*_*k*_ distances, it is important to keep in mind that the adjacency spectral distance compares the *largest k* eigenvalues, whereas the Laplacian spectral distances compare the *smallest k* eigenvalues. Comparison using the first *k* eigenvalues for small *k* allows one to focus on the community structure of the graph, while ignoring the local structure of the graph [15]. Inclusion of the higher-*k* eigenvalues allows one to discern local features as well as global. As we will see, this flexibility allows the user to target the particular scale at which they wish to examine the graph, and is a significant advantage of the spectral distances.

The three spectral distances used here are not true metrics. This is because there exist graphs *G* and *G′* that are co-spectral but not isomorphic. That is to say, adjacency cospectrality occurs when 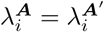 for all *i* = 1, *…, n*, so *d*_***A***_(*G, G′*) = 0, but *G* ≇*G′*. Similar notions of cospectrality exist for all matrix representations; graphs that are co-spectral with respect to one matrix representation are not necessarily co-spectral with respect to other representations.

Little is known about cospectrality, save for some computational results on small graphs [16] and trees [13]. Schwenk proved that a sufficiently large tree nearly always has a co-spectral counterpart [17]. This result was extended recently to include a wide variety of random trees [18]. However, results such as these are not of great import to us; the graphs examined are large enough that we do not encounter cospectrality in our numerical experiments. A more troubling failure mode of the spectral distances would be when the distance between two graphs is very small, but the two graphs have important topological distinctions. In Section, we will provide further insight into the effect of topological changes on the spectra of some of the random graph models we study.

The consideration above addresses the question of how local changes effect the overall spectral properties of a graph. Some limited computational studies have been done in this direction. For example, Farkas et al. [19] study the transition of the adjacency spectrum of a small world graph as the disorder parameter increases. As one might expect, they observe the spectral density transition from a highly discontinuous density (which occurs when the disorder is zero, and so the graph is a ring-like lattice) to Wigner’s famous semi-circular shape [20] (which occurs when the disorder is maximized, so that the graph is roughly equivalent to an uncorrelated random graph.)

From an analytical standpoint, certain results in random matrix theory inform our understanding of fluctuations of eigenvalues of the uncorrelated random graph (see Section for a definition). These results hold asymptotically as we consider the *k*^th^ eigenvalue of a graph of size *n*, where *k* = *αn* for *α ∈* (0, 1]. In this case, O’Rourke [21] has shown that the the eigenvalue *λ*_*k*_ is asymptotically normal with asymptotic variance *σ*^2^(*λ*_*k*_) = *C*(*α*) log *n/n*. An expression for the constant *C*(*α*) is provided; see Remark 8 in [21] for the detailed statement of the theorem. This result can provide a heuristic for spectral fluctuations in some random graphs, but when the structure of these graphs diverges significantly from that of the uncorrelated random graph, then results such as these become less informative.

Another common question is that of interpretation of the spectrum of a given matrix representation of a graph.^4^ How are we to understand the shape of the empirical distribution of eigenvalues? Can we interpret the eigenvalues which separate from this bulk in a meaningful way? The answer to this question depends, of course, on the matrix representation in question. Let us focus first on the Laplacian matrix ***L***, the interpretation of which is the clearest.

The first eigenvalue of ***L*** is always *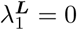*, with the eigenvector being the vector of all ones, **1** *∈* ℝ^*n*^. It is a well-known result that the multiplicity of the zero eigenvalue is the number of connected components of the graph, i.e. if 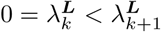, then there are precisely *k* connected components of the graph [22]. Furthermore, in such a case, the first *k* eigenvectors will indicate the components. In [15], an approximate version of this statement is made rigorous, in which the first *k* eigenvalues being small is an indicator of a graph being strongly partitioned into *k* clusters. This result justifies the use of the Laplacian in spectral clustering algorithms.

The eigenvalues of the Laplacian also have an interpretation analogous to the vibrational frequencies that arise as the eigenvalues of the continuous Laplacian operator *∇*^2^. To understand this analogy, consider the graph as embedded in a plane, with each vertex representing an oscillator of mass one and each edge a spring with elasticity one. Then, for small oscillations perpendicular to the plane, the Laplacian matrix is precisely the coupling matrix for this <u>sys</u>tem, and so the eigenvalues give the square of the normal mode frequencies, 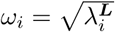. For a more thorough exposition of this interpretation of the Laplacian, see [23].

Maas [24] suggests a similar interpretation of the spectrum of the adjacency matrix ***A***. Consider the graph as a network of oscillators, embedded in a plane as previous. Additionally suppose that each vertex is connected to so many external non-moving points (by edges with elasticity one) so that the graph becomes regular with degree *r*. The frequencies of the normal modes of this structure then connect to the eigenvalues of ***A*** via 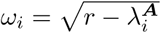.^5^

#### Matrix Distances

The second class of distances we will discuss are called *matrix distances*, and consist of direct comparison of the structure of pairwise affinities between vertices in a graph. These affinities are frequently organized into matrices, and the matrices can then be compared, often via an entry-wise 𝓁_*p*_ norm.

We have discussed spectral methods for measuring distances *between* two graphs; to introduce the matrix distances, we will begin by focusing on methods for measuring distances *on graphs*; that is to say, the distance *δ*(*v, w*) between two vertices *v, w ∈ V*. Just a few examples of such distances include the shortest path distance [25], the effective graph resistance [26], and variations on random-walk distances [27]. Of those listed above, the shortest path distance is the oldest and the most thoroughly studied; in fact, it is so ubiquitous that “graph distance” is frequently used synonymously with shortest path distance [28].

There are important differences between the distances *δ* that we might choose. The shortest path distance considers only a single path between two vertices. In comparison, the effective graph resistance takes into account all possible paths between the vertices, and so measures not only the length, but the *robustness* of the communication between the vertices. This distinction is important when, for example, considering travel between two locations on a road network subject to high traffic.

How do these distances *on a graph* help us compute distances *between graphs*? Let us denote by *δ* : *V × V* → ℝ a generic distance on a graph. We need assume very little about this function, besides it being real-valued; in particular, it need not be symmetric, and we can even allow *δ*(*v, v*) ≠ 0.^6^ Recalling that our vertices *v ∈ V* = *{*1, *…, n}* are labelled with natural numbers, we can then construct a matrix of pairwise distances ***M*** via 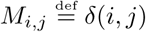.

The idea behind what we refer to as **matrix distances** is that this matrix ***M*** carries important structural information about the graph. Suppose that, for our given distance *δ*(·*,·*) graphs *G* and *G′* have corresponding matrices ***M*** and ***M*** *′*. We can then compare *G* and *G′* via

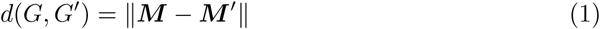

where ‖ · ‖ is a norm we are free to choose.^7^

Let us elucidate a specific example of such a distance; in particular, we will show how the edit distance conforms to this description. Let *δ*(*v, w*) be defined as

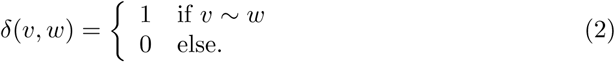

Then the matrix ***M*** is just the adjacency matrix ***A***. If we use the norm

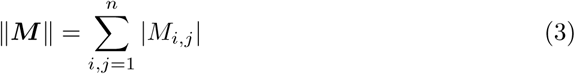

then we call the resulting distance 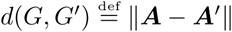 the **edit distance**.

Of course, the usefulness of such a distance is directly dependent on how well the matrix ***M*** reflects the topological structure of the graph. The edit distance focuses by definition on local structure; it can only see changes at the level of edge perturbations. If significant volume changes are happening in the graph, then the edit distance will detect this, as do our other matrix distances. However, in our numerical experiments, we match the expected volume of the models under comparison, and so the edit distance is generally unable to discern between the models seen in Section.

We also implement the resistance-perturbation distance, first discussed in [11]. This distance takes the effective graph resistance *R*(*u, v*), defined in [26], as the measure of vertex affinity. This results in a (symmetric) matrix of pairwise resistances ***R***. The resistance-perturbation distance (or just resistance distance) is based on comparing these two matrices in the entry-wise 𝓁_1_ norm given in Equation (3).

The nice theoretical properties of the effective graph resistance [26] motivate our computational exploration of how well it reflects structure in realistic scenarios. Unlike the edit distance, the resistance distance is designed to detect global structural differences between graphs. A recent work [29] discusses the efficacy of the resistance distance in detecting community changes.

Finally, we look at DELTACON, a distance based on the fast belief-propagation method of measuring node affinities [5]. To compare graphs, this method uses the fast belief-propagation matrix

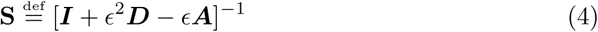

and compares the two representations **S** and **S***′* via the Matusita difference:

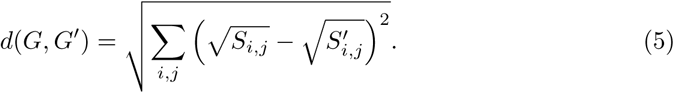

Note that the matrix **S** can be rewritten in a matrix power series as

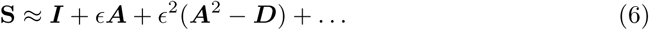

and so takes into account the influence of neighboring vertices in a weighted manner, where neighbors separated by paths of length *k* have weight *ϵ*^*k*^. Fast belief-propagation is designed to model the diffusion of information throughout a graph [30], and so should in theory be able to perceive both global and local structures. Although empirical tests are performed in [5], no direct comparison to other modern methods are presented.

#### Other Graph Distances

These two categories do not cover all possible methods of graph comparison. The computer science literature explores various other methods (see [8], Section 3.2 for a nice review), and other disciplines that apply graph-based techniques often have their own idiosyncratic methods for comparing graphs extracted from data.

One possible method for comparing graphs is to look at specific “features” of the graph, such as the degree distribution, betweenness centrality distribution, diameter, number of triangles, number of *k*-cliques, etc. For graph features that are vector-valued (such as degree distribution) one might also consider the vector as an empirical distribution and take as graph features the sample moments (or quantiles, or statistical properties). A **feature-based distance** is a distance that uses comparison of such features to compare graphs.

Of course, in a general sense, all methods discussed so far are feature based; however, in the special case that the features occur as values over the space *V × V* of possible node pairings, we choose to refer to them more specifically as matrix distances. Similarly, if the feature in question is the spectrum of a particular matrix realization of the graph, we will call the method a spectral distance.

In [31], a feature-based distance called NetSimile is proposed, which focuses on local and egonet-based features (e.g. degree, volume of egonet as fraction of maximum possible volume, etc). If we are using *k* features, the method aggregates a feature-vertex matrix of size *k × n*. This feature matrix is then reduced to a “signature vector” (a process they call “aggregation”) which consists of the mean, median, standard deviation, skewness, and kurtosis of each feature. These signature vectors are then compared in order to obtain a measure of distance between graphs.

In the neuroscience literature in particular, feature-based methods for comparing graphs are popular. In [32], the authors use graph features such as modularity, shortest path distance, clustering coefficient, and global efficiency to compare functionally connectivity networks of patients with and without schizophrenia. Statistics of these features for the control and experiment groups are aggregated and compared using standard statistical techniques.

We implement NETSIMILE in our numerical tests as a prototypical feature-based method. It is worth noting that the general approach could be extended in almost any direction; any number of features could be used (which could take on scalar, vector, or matrix values) and the aggregation step can include or omit any number of summary statistics on the features, or can be omitted entirely. We implement the method as it is originally proposed, with the caveat that calculation of many of these features is not appropriate for large graphs, as they cannot be computed in linear or near-linear time. A scalable modification of NETSIMILE would utilize features that can be calculated (at least approximately) in linear or near-linear time.

#### Scaling and Complexity of Algorithms

In many interesting graph analysis scenarios, the sizes of the graphs to be analyzed are on the order of millions or even billions of vertices. For example, the social network defined by Facebook users has over 2 billion vertices as of 2017. In scenarios such as these, any algorithm of complexity 𝒪(*n*^2^) will become unfeasible; although in principle it is possible that the constant *M* would be so small it would make up for the *n*^2^ term in the complexity, in practice this is not the case. This motivates our requirement that our algorithms be of near-linear complexity. Indeed, even for graphs on the scale of 10^5^, quadratic algorithms quickly become unfeasible.

This challenge motivates the previously stated requirement that all algorithms be of linear or near-linear complexity. We say an algorithm is **linear** if it is 𝒪(*n*); it is **near-linear** if it is 𝒪 (*n* log *a*_*n*_) where *a*_*n*_ is asymptotically bounded by a polynomial. We use the notation *a*_*n*_ = 𝒪 (*b*_*n*_) in the standard way; for a more thorough discussion of algorithmic complexity, including definitions of the 𝒪 notation, see [33].

We focus our attention on sparse graphs. We define sparsity as an asymptotic property, and so it is only defined on a sequence of graphs. However, one can reasonably apply this to empirical graph data which changes over time and thus generates a natural time series which can be tested (roughly, since we are always at finite time) against this definition. In particular, let 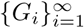 be a sequence of graphs, where the size of *G*_*i*_ is *i* and the number of edges in *G*_*i*_ denoted by *m*_*i*_. We say a graph is sparse when the sequence 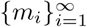 is near-linear, in the sense given above.

Table 1 indicates the algorithmic complexity of each distance measure we compare. For DELTACON and the resistance distance, there are approximate algorithms as well as exact algorithms; we list the complexity of both. Although we use the exact versions in our experiments, in practice the approximate version would likely be used if the graphs to be compared are large.

**Table 1.** Distance measures and complexity. *n* indicates the maximum of size of the two graphs being compares, and *m* indicates the maximum number of edges. For the spectral decomposition, *k* denotes the number of principal eigenvalues we wish to find. We assume that factors such as graph weights and quality of approximation are held constant, leading to simpler expressions here than appear in cited references. Spectral distances have equivalent complexity, since they all all amount to performing an eigendecomposition on a symmetric real matrix.

| Distance Measure | Complexity | Ref. |
| --- | --- | --- |
| Edit Distance | $\mathcal{O}(m)$ | 8 |
| Resistance Distance (Exact) | $\mathcal{O}(n^2)$ | [11] |
| Resistance Distance (Approximate) | $\mathcal{O}(m)$ | [11] |
| DeltaCon (Exact) | $\mathcal{O}(n^2)$ | [5] |
| DeltaCon (Approximate) | $\mathcal{O}(m)$ | [5] |
| NetSimile | $\mathcal{O}(n \log n)$ | [31] |
| Spectral Distance | $\mathcal{O}(nk^2)$ | [34] |

Of particular interest are the highly parallelizable randomized algorithms which can allow for extremely efficient matrix decomposition. In [34], the authors review many such algorithms, and discuss in particular their applicability to determining principal eigenvalues. The computation complexity in Table 1 for the spectral distances is based on their simplified analysis of the Krylov subspace methods, which states that the approach is 𝒪 (*kT*_mult_ + (*m* + *n*)*k*^2^), where *T*_mult_ is the cost of matrix-vector multiplication for the input matrix. Since our input matrices are sparse, *T*_mult_ = 𝒪 (*n*), and *m* + *n* = 𝒪 (*n*). Although we use the implicitly restarted Arnoldi method in our eigenvalue calculations, if implementing such a decomposition on large matrices the use of a randomized algorithm could lead to a significant increase in efficiency.

### Random Graph Models

Random graph models have long been used as a method for understanding topological properties of graph data that occurs in the world. The uncorrelated random graph model of Erdős and Rényi [12] is the simplest model, and provides a null model akin to white noise. The tractability of this model has led to some beautiful probabilistic analysis [35] but the uniform topology of the model does not accurately model empirical graph data. The stochastic blockmodel is an extension of the uncorrelated random graph, but with explicit community structure reflected in the distribution over edges.

Models such as preferential attachment [9] and the Watts-Strogatz model [10] have been designed to mimic properties of observed graphs. Very little can be said about these models analytically, and thus much of what is understood about them is computational. The two-dimensional square lattice is a quintessential example of a highly structured and regular graph.

We will now introduce the models that are used. We study only undirected graphs, with no self-loops. Although directed graphs of are of great practical importance [36], tools such as the graph resistance only apply to undirected graphs. In particular, the electrical analogy needed to render the effective graph resistance meaningful is lost in a directed graph. Random-walk concepts are still perfectly meaningful on directed graphs, and motivate many popular algorithms used on such graphs (see e.g. [37]).

Most of the models in this work are sampled via the Python package NETWORKX [38]; details of implementation can be found in the source code of the same. Some of the models we use are most clearly defined via their associated probability distribution, while others are best described by a generative mechanism. We will introduce the models roughly in order of complexity.

#### The Uncorrelated Random Graph

The **uncorrelated random graph** (also known as the Erdős-Rényi random graph) is a random graph in which each edge exists with probability *p*, independent of the existence of all others. We denote this distribution of graphs by *G*(*n, p*) (recall that *n* denotes the size of the graph). As previously mentioned, this is by far the most thoroughly studied of random graph models; the simplicity of its definition allows for analytic tractability not found in many other models of interest. For example, the spectrum of the uncorrelated random graph is well understood. In particular, the spectral density forms a semi-circular shape, first described by Wigner [20], of radius 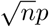, albeit with a single eigenvalue 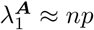 separate from the semicircular bulk [19].

We will employ the uncorrelated random graph as our null model in many of our experiments. It is, in some sense, a “structureless” model; more specifically, the statistical properties of each edge and vertex in the graph are exactly the same. This model fails to produce many of properties observed in empirical networks, which motivates the use of alternative graph models. For a more detailed definition of the model, and a thorough study of the properties of the uncorrelated random graph, see [35].

#### The Stochastic Blockmodel

One important property of empirical graphs is community structure. Vertices often form (relatively) densely connected communities, with the connection between communities being (relatively) sparse, or non-existent. This motivates the use of the **stochastic blockmodel**. In this model, each of the *n* vertices are in one two non-overlapping sets *C*_1_ and *C*_2_, referred to as “communities”. Each edge *e* = (*i, j*) exists (independently) with probability *p* if *i* and *j* are in the same community, and *q* if *i* and *j* are in separate communities. In this work, we will use “balanced” communities, so that the difference in size |*C*_1_| *–* |*C*_2_| is less than 1 in magnitude.

The stochastic blockmodel is a prime example of a model which exhibits global structure without any meaningful local structure. In this case, the global structure is the partitioned nature of the graph as a whole. On a file scale, the graph looks like an uncorrelated random graph. We will use the model to determine which distances are most effective at discerning global (and in particular, community) structure.

The stochastic blockmodel is at the cutting edge of rigorous probabilistic analysis of random graphs. In particular, Abbe et al. [39] have recently proven a strict bound on community recovery, showing in exactly what regimes of *p* and *q* it is possible to discern the communities.

Generalizations of this model exist in which there are *K* communities of arbitrary size. Furthermore, each community need not have the same parameter *p*, and each community *pair* need not have the same parameter *q*. One can, in general, construct a (symmetric) matrix of parameters with the *p*_*i*_ on the diagonal, and the *q*_*i,j*_ elsewhere, for 1 *≤ i, j ≤ K*. However, since our model has only two communities of nearly equal size, we only need a pair of parameters (*p, q*).

#### Preferential Attachment Models

Another often-studied feature of empirical graphs are their degree distribution. This is generally visualized as a histogram of degree frequencies, and studied under the assumption that it reflect some underlying distribution that informs us about the generative mechanism of the graph.

The degree distribution of an uncorrelated random graph is binomial, and so it has tails that exponentially decrease; for large *d*, the probability P[*d*] that a randomly chosen vertex has degree *d* decays exponentially, ℙ [*d*] *∝* exp(*-d*^2^). However, in observed graphs such as computer networks, human neural nets, and social networks, the observed degree distribution has a power-law tail [9]. In particular, one observed ℙ[*d*] *∝ d*^*-γ*^ where generally *γ ∈* [2, 3]. Such distributions are often also referred to as “scale-free”.

The **preferential attachment model** is a scale-free random graph model. It is best described via the generative process rather than by a particular distribution over edges or possible graphs.^9^ Although first described by Yule in 1925 [40], the model did not achieve its current popularity until the work of Barabási and Albert in 1999 [9].

The model has two parameters, *l* and *n*. The latter is the size of the graph, and the former controls the density of the graph. We require that 1 ≤ *l < n*. The generative procedure for sampling from this distribution proceeds as follows. Begin by initializing a star graph with *l* + 1 vertices, with vertex *l* + 1 having degree *l* and all others having degree 1. Then, for each *l* + 1 *< i* ≤ *n*, add a vertex, and randomly attach it to *l* vertices already present in the graph, where the probability of *i* attaching to *v* is proportional to to the degree of *v*. Stop once the graph contains *n* vertices.

The constructive description of the algorithm does not yield itself to simple analysis, and so less is known analytically about the preferential attachment model than the uncorrelated random graph or the stochastic blockmodel. In [41], the authors prove that the *k*^th^ eigenvalue of the Laplacian 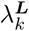 scales like 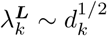, where *d*_*k*_ is the *k*^th^ largest degree in the graph.^10^ In [19], the authors demonstrate numerically that the adjacency spectrum exhibits a triangular peak with power-law tails.

Having a high degree makes a vertex more likely to attract more connections, so the graph quickly develops strongly connected “hubs,” or vertices of high degree. This impacts both the global and local structure of the graph. Hubs are by definition global structures, as they touch a significant portion of the rest of the graph, making path lengths shorter and increasing connectivity throughout the graph. On the local scale, vertices in the graph tend to connect exclusively to the highest-degree vertices in the graph, rather than to one another, generating a tree-like topology. This particular topology yields a signature in the tail of the spectrum, the importance of which will be discussed below.

#### The Watts-Strogatz Model

Many real-world graphs exhibit the so-called “small world pheomenon,” where the expected shortest path length between two vertices chosen uniformly at random grows logarithmically with the size of the graph. Watts and Strogatz [10] constructed a random graph model that exhibits this behavior, along with a high clustering coefficient not seen in an uncorrelated random graph. The clustering coefficient is defined as the ratio of number of triangles to the number of connected triplets of vertices in the graph. The **Watts-Strogatz model** [10] is designed to be the simplest random graph that has high local clustering and small average (shortest path) distance between vertices.

Like preferential attachment, this graph is most easily described via a generative mechanism. The algorithm proceeds as follows. Let *n* be the size of the desired graph, let 0 *≤ p ≤* 1, and let *k* be an even integer, with *k < n*. We begin with a ring lattice, which is a graph where each vertex is attached to its *k* nearest neighbors, *k/*2 on each side. Then, for each edge in the graph (*i, j*) with *i < j*, with probability *p* rewire the edge to a random vertex *l*, so that (*i, j*) is replaced with (*i, l*). The target *l* is chosen so that *i* ≠ *l* and *i* ≁*l* at the time of rewiring. Stop once all edges have been iterated through. In our implementations, we add an additional stipulation that the graph must be connected. If the algorithm terminates with a disconnected graph, then we restart the algorithm and generate a new graph. This process is repeated until the resulting graph is connected.

As mentioned before, the topological features that are significant in this graph are the high local clustering and short expected distance between vertices. Of course, these quantities are dependent on the parameter *p*; as *p* **→** 1, the Watts-Strogatz model approaches an uncorrelated random graph. Similarly, as *p* **→** 0 the adjacency spectral density transitions from the tangle of sharp maxima typical of a ring-lattice graph to the smooth semi-circle of the uncorrelated random graph [19]. Unlike the models above, this model exhibits primarily local structure. Indeed, we will see that the most significant differences lie in the tail of the adjacency spectrum, which can be directly linked to the number of triangles in the graph [19]. On the large scale, however, this graph looks much like the uncorrelated random graph, in that it exhibits no communities or high-degree vertices.

This model fails to produce the scale-free behavior observed in many empirical graph data sets. Although the preferential attachment model reproduces this scale-free behavior, it fails to reproduce the high local clustering that is frequently observed, and so we should think of neither model as fully replicating the properties of observed graphs.

#### Random Degree-Distribution Graphs

The above three models are designed to mimic certain properties of empirical graphs. In some cases, however, we may wish to fully replicate a given degree sequence, while allowing other aspects of the graph to remain random. That is to say, we seek a distribution that assigns equal probability to each graph, conditioned upon the graph having a given degree sequence. The simplest model that attains this result is the configuration model [42]. Recently, Zhang et al. [43] have derived an asymptotic expression for the adjacency spectrum of a configuration model, which is exact in the limit of large graph size and large mean degree.

Inconveniently, this model is not guaranteed to generate a simple graph; the resulting graph can have self-edges, or multiple edges between two vertices. In 2010, Bayati et al. [44] described an algorithm which uniformly samples from the space of simple graphs with a given degree distribution. We will refer to graphs sampled in this way as **random degree-sequence graphs**. Their utility lies in the fact that we can use them to control for the degree sequence when comparing graphs; they are used as a null model, similar to the uncorrelated random graph, but they can be tuned to share some structure (notably, the power-law degree distribution of preferential attachment) with the graphs to which they are compared.

The generative algorithm for this model is designed to sample from a uniform distribution over all possible graphs of a given size, conditioned upon the provided degree distribution. Their algorithm is fast, but not perfectly uniform; in [44] the authors prove that the distribution is asymptotically uniform, but do not prove results for finite graph size. We use this algorithm despite the fact that it does not sample the desired distribution in a truly uniform manner; the fact that the resulting graph is simple overcomes this drawback.

#### Lattice Graphs

In some of our experiments, we utilize lattice graphs. In particular we use a 2-dimensional *n* by *m* rectangular lattice. Using such a predictable structure allows us to test our understanding of our distances; in particular, we can see if our distances respond as we expect to structural features that are present in the lattice. Empirical realizations of planar graphs, such as road networks, often exhibit lattice-like structure. The planar structure of the lattice allows for an intuitive understanding of the spectral features of the graph, as they approximate the normal vibrational frequencies of a two-dimensional surface.

Lattice graphs are highly regular, in the sense that the connectivity pattern of each (interior) vertex is identical. This regularity is reflected by the discrete nature of the lattice’s spectrum, which can be seen in Figure 10. This is a particularly strong flavor of local structure, as it is not subject to the nose present in random graph models. This aspect allows us to probe the functioning of our distances when they are exposed to graphs with a high amount of inherent structure and very low noise.

**Fig 1.**
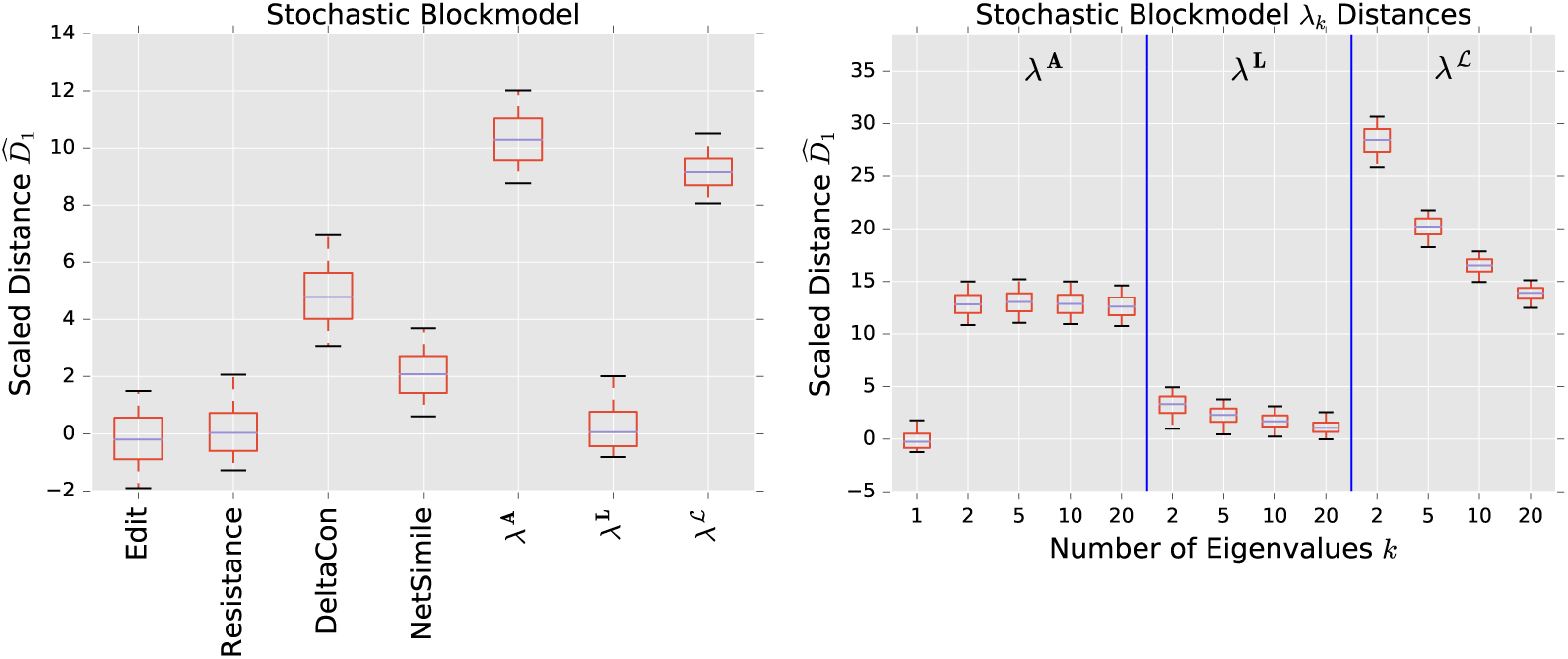
Comparison of distance performance, with uncorrelated random graph as null model and stochastic blockmodel as alternative. Boxes extend from lower to upper quartile, with center line at median. Whiskers extend from 5th to 95th percentile.

**Fig 2.**
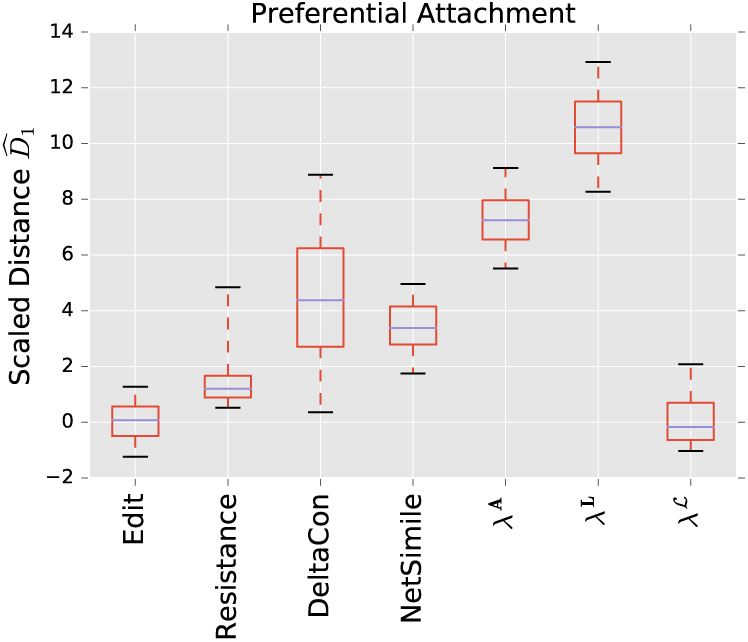
Comparison of distance performance, with uncorrelated random graph as null model and preferential attachment as alternative. See Figure 1 for boxplot details.

**Fig 3.**
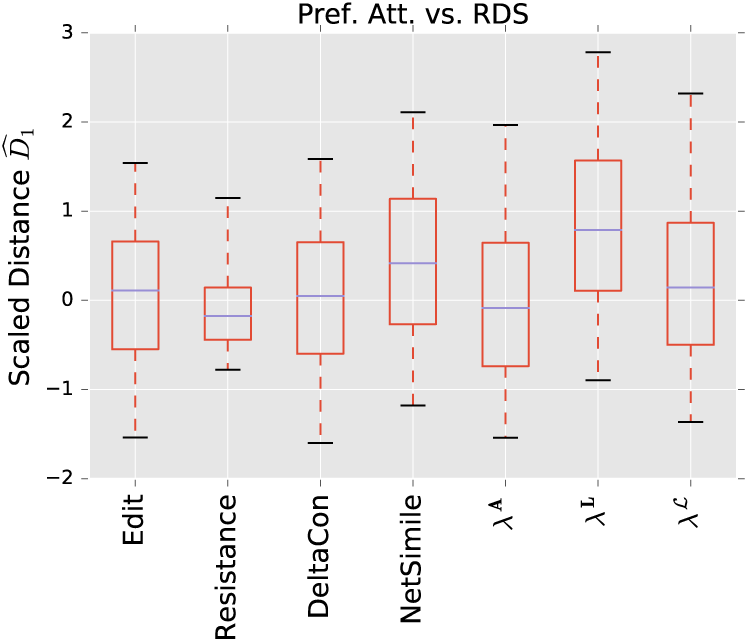
Comparison of distance performance, with degree-sequence random graph as null model and preferential attachment as alternative. The degree sequence for each null matches that of the alternative. See Figure 1 for boxplot details.

**Fig 4.**
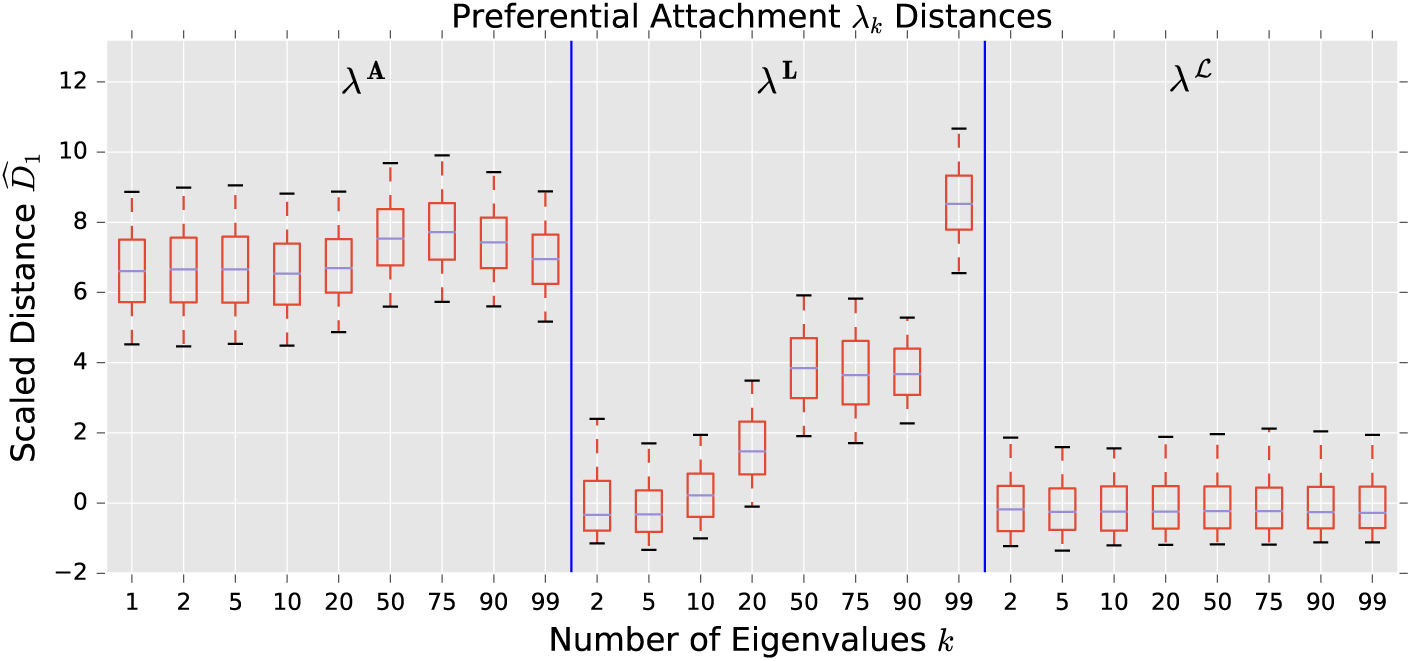
Comparison of *λ*_*k*_ distance performance, with uncorrelated random graph as null model and preferential attachment as alternative. See Figure 1 for boxplot details.

**Fig 5.**
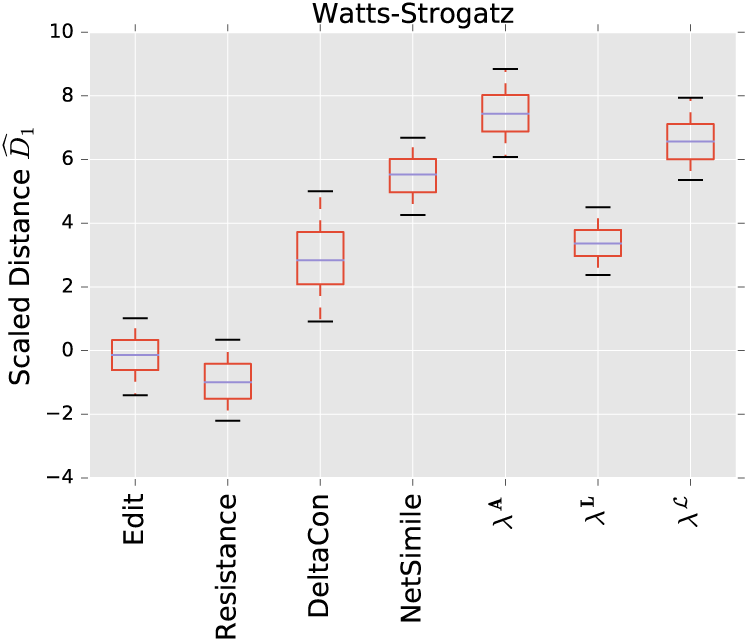
Comparison of distance performance, with uncorrelated random graph as the null model, and a small-world graph as the alternative. See Figure 1 for boxplot details.

**Fig 6.**
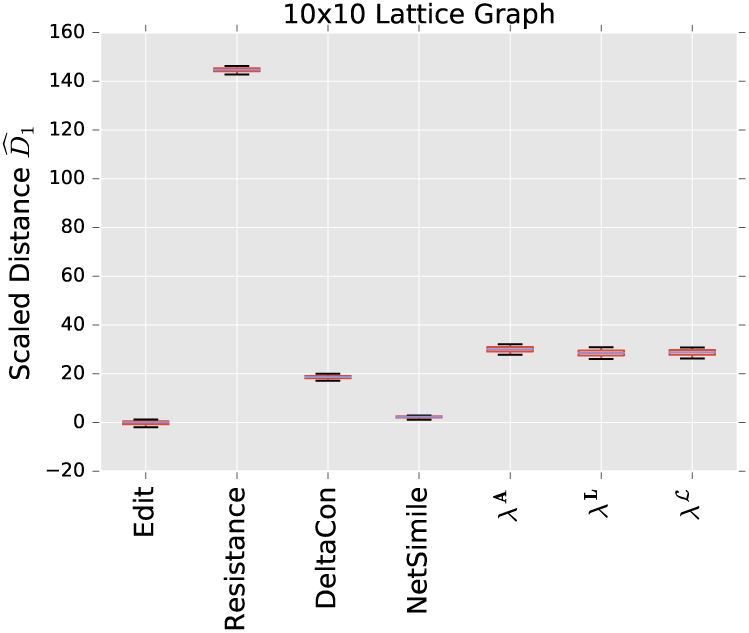
Comparison of distance performance, with a 10 by 10 2-dimensional lattice graph as the alternative model, and a random degree-distribution graph (with the same degree distribution as the lattice) as the null. See Figure 1 for boxplot details.

**Fig 7.**
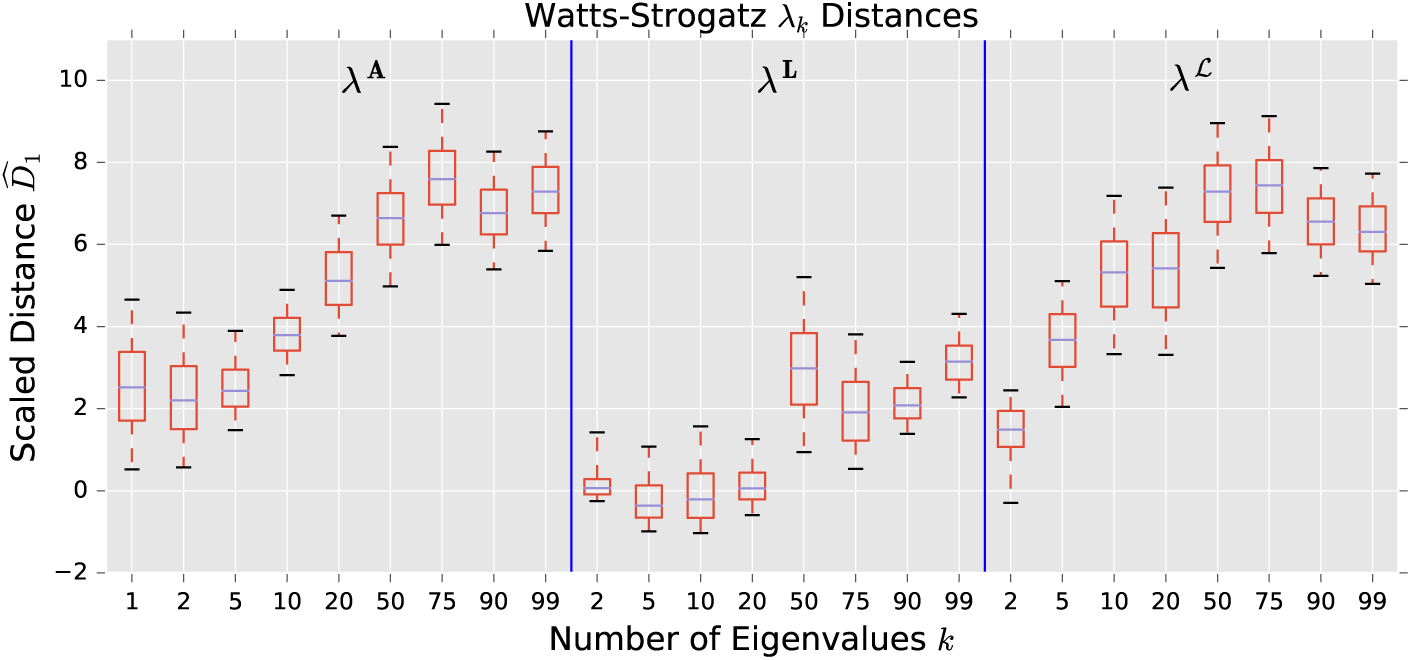
Comparison of *λ*_*k*_ distance performance, with uncorrelated random graph as the null model, and a small-world graph as the alternative. See Figure 1 for boxplot details.

**Fig 8.**
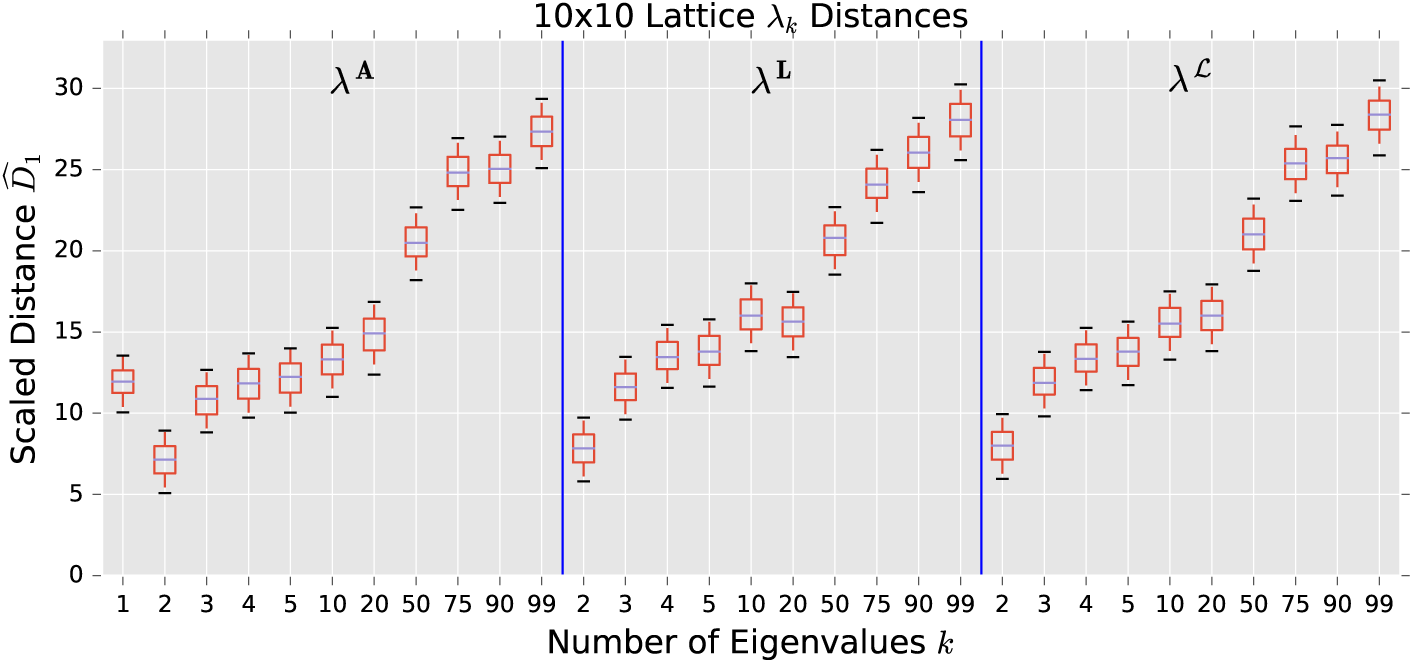
Comparison of *λ*_*k*_ distance performance, with a 10 by 10 2-dimensional lattice graph as the alternative model, and a random degree-distribution graph (with the same degree distribution as the lattice) as the null. See Figure 1 for boxplot details.

**Fig 9.**
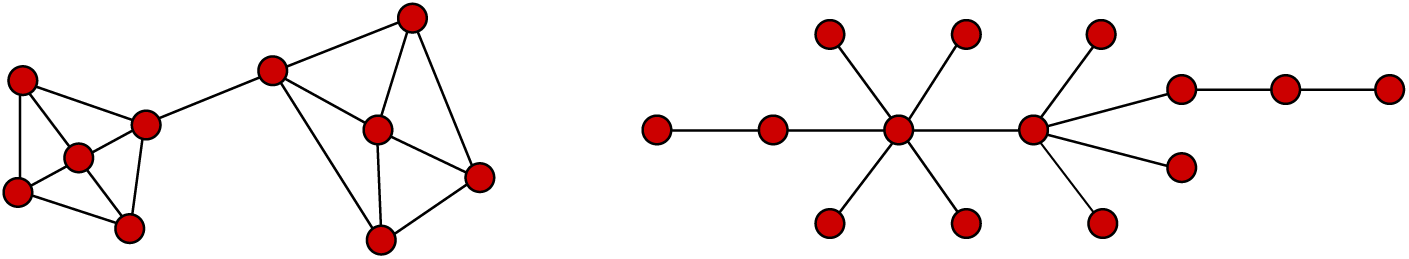
Two significant global structures observed in our experiments. On the left is the community structure typical of the stochastic blockmodel. On the right is the heavy-tailed degree distribution typical of the preferential attachment model.

**Fig 10.**
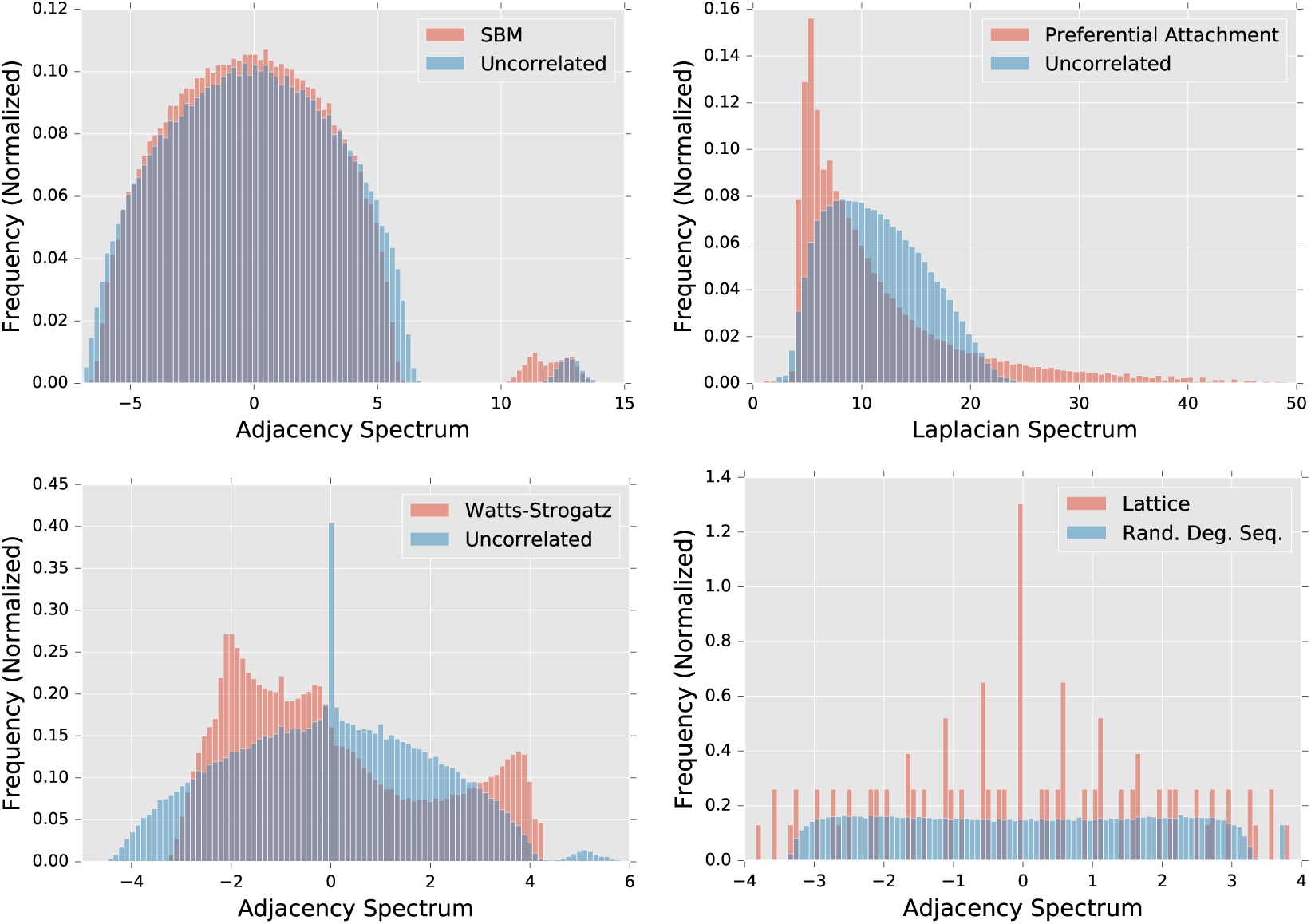
Spectral densities for various graph comparisons. Parameters used match those in Sections through. Densities are built from an ensemble of 1000 graphs. The Laplacian spectrum *λ*^***L***^ is shown for preferential attachment, while adjacency spectrum *λ*^***A***^ is shown for all others. The uncorrelated random graph model in the lower left has a lower *p* than those on the upper row, resulting in a sharp peak at *λ*^***A***^ = 0.

#### Exponential Random Graph Models

A popular random graph model is the **exponential random graph model**, or ERGM for short. Although they are popular and enjoy simple interpretability, we do not use ERGMs in our experiments. Unlike some of our other models which are described by their generative mechanisms, these are described directly via the probability of observing a given graph *G*.

In particular, let *g*_*i*_(*G*) be some scalar graph properties (e.g. size, volume, or number of triangles) and let *θ*_*i*_ be corresponding coefficients, for *i* = 1, *…, K*. Then, the ERGM assigns to each graph a probability [45]

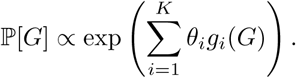

This distribution can be sampled via a Gibbs sampling technique, a process which is outlined in detail in [45]. ERGMs show great promise in terms of flexibility and interpretability; one can seemingly tune the distribution towards or away from any given graph metric, including mean clustering, average path length, or even decay of the degree distribution.

However, our experience attempting to utilize ERGMs led us away from this approach. When sampling from ERGMs, we were unable to control properties individually to our satisfaction. In particular, we found that attempts to increase the number of triangles in a graph increased the graph volume; when we subsequently used the ERGM parameters to *de-emphasize* graph volume, the sampled graphs had an empirical distribution very similar to an uncorrelated random graph.

### Evaluation of Distances on Random Graph Ensembles

We will now present the results of our numerical tests, which compare the effectiveness of the various distances in discerning between pairs of random graph ensembles. The discussion in this section will be brief; we reserve our interpretation of the results until the next section. The experiments are organized via the models being compared, and with the performance of each distance shown in plots in each section. When appropriate, we also show the performance of the *λ*_*k*_ distances for various *k*. Table 2 summarizes each comparison performed.

**Table 2.** Table of comparisons performed, and the important structural features therein. *G*(*n, p*) indicates the uncorrelated random graph, SBM is the stochastic blockmodel, PA is the preferential attachment model, RDDG is the random degree distribution graph, and WS is the Watts-Strogatz model.

| Section | Null | Alternative | Primary Structural Difference |
| --- | --- | --- | --- |
| | $G(n, p)$ | SBM | Community structure |
| | $G(n, p)$ | PA | High-degree vertices |
|  | RDDG | PA | Structure not in degree distr. |
| | $G(n, p)$ | WS | Local structure |
| | $G(n, p)$ | Lattice | Extreme local structure |

#### Description of Experiments

The experiments are designed to mimic a scenario in which a practitioner is attempting to determine whether a given graph belongs to a population or is an outlier relative to that population. In this vein, we perform experiments that determine how well each distance distinguishes populations drawn from a random graph model. In particular, let us define by 𝒢_0_ and 𝒢_1_ our two graph populations, which we will refer to as the **null** and **alternative** populations, respectively. For each distance measure, let 𝒟_0_ be the distribution of distances 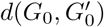 where *G*_0_ and 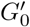 are both drawn from the distribution ***𝒢***_0_. Similarly, let ***𝒟***_1_ be the distribution of distances *d*(*G*_0_, *G*_1_), where *G*_0_ is drawn from ***𝒢***_0_ and *G*_1_ is drawn from ***𝒢***_1_.

The distances ***𝒟***_0_ are a characteristic distance between members of the population ***𝒢***_0_. The intuition here is that if the distribution of ***𝒟***_1_ is well separated from that of ***𝒟***_0_, then that distance is effective at separating the null population from the alternative population; if member of ***𝒢***_1_ are much further from members of ***𝒢***_0_ than this characteristic in-population distance, we can easily distinguish the two.

To that end, we normalize the statistics of ***𝒟***_1_ by those of ***𝒟***_0_ in order to compare. In particular, let *µ*_*i*_ be the sample mean of ***𝒟***_*i*_, and let *σ*_*i*_ be the sample standard deviation, for *i ∈ {* 0, 1 *}*. Then, we examine the statistics of 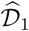, whose samples 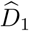 are calculated via

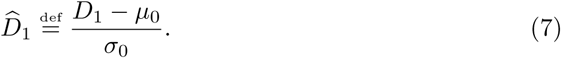

If our empirical distribution of 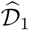 is well separated from zero (i.e. the mean is significantly greater than the standard deviation), then the distance is effectively separating the null and alternative population.

We generate 500 samples of *D*_0_ and *D*_1_, where each sample compares two graphs of size *n* = 100, unless otherwise specified. The graphs are always connected; our sampler will discard a draw from a random graph distribution if the resulting graph is disconnected. Said another way, we draw from the distribution defined by the model, conditioning upon the fact that the graph is connected.

The small size of our graphs allows us to use larger sample sizes; although all of the matrix distances used have fast approximate algorithms available, we use the slower, often 𝒪(*n*^2^), exact algorithms for our experiments, and so larger graphs would be prohibitively slow to work with. In all the below experiments, we choose our parameters so that the expected volume of the two models under comparison is equal.

Sections through are separated by the models being compared. A very brief discussion of the results occurs alongside the presentation of the results, while a more thorough discussion is reserved for Section. The reader who wishes to primarily understand our observations and interpretation of the results can skip to and reference the above sections as necessary.

## Experimental Results

### Stochastic Blockmodel

In Figure 1, we see the results of comparison between an uncorrelated random graph model and a stochastic block model. For the uncorrelated random graph, the probability of an edge existing is *p* = 0.12, which is a value of *p* for which the graph was almost always connected.^11^ For the stochastic blockmodel, we have two communities, each of size *n/*2 = 50, with parameters (*p, q*) = (0.228, 0.012). Thus, the in-community connectivity is more dense than the cross-community connectivity by a factor of *p/q* = 19.

Since we have volume matched the graphs, the edit distance fails to distinguish the two models. Among the matrix distances, DELTACON separates the two models most reliably. The adjacency and normalized Laplacian distances perform well, but the non-normalized Laplacian distance fails to distinguish the two models. The performance of the adjacency distance is primarily in the second eigenvalue 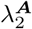, and including further eigenvalues adds no benefit; the normalized Laplacian also shows most of its benefit in the second eigenvalue 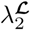, but unlike the adjacency distance, including more eigenvalues decreases the performance of the metric.

### Preferential Attachment vs Uncorrelated

Figure 2 shows the results of comparing a preferential attachment graph to an uncorrelated random graph. The preferential attachment graph is quite dense, with *l* = 6. Since the number of edges in this model is always |*E*| = *l*(*n – l*), we calculate the parameter *p* for the uncorrelated graph via

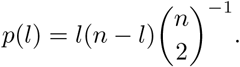

We use *p*(6) = 0.12 in these experiments.

Again, due to matching the volumes of the graph, the edit distance fails to distinguish the two models. The resistance distance shows mediocre performance, although 0 is outside the 95% confidence interval. DeltaCon exhibits extremely high variability, although it has the highest median of the matrix distances.

The Laplacian distance outperforms all others, while the normalized Laplacian does not separate the two models at all. Figure 4 shows that most of the spectral information for the Laplacian is contained in the last few eigenvalues, counter to what one often expects from *λ*_*k*_ distances. For the adjacency distance, most of the information is held in the first eigenvalue, as the scaled distance stays more or less constant as one increases *k* (plot not shown).

### Preferential Attachment vs Random Degree Distribution Graph

In addition to the comparison of preferential attachment and uncorrelated random graphs in Section, we now compare preferential attachment to random degree-distribution graphs. Recall that for a given degree distribution, the random degree-distribution graph probability density is the uniform density over all **simple** graphs with the given degree distribution. We employ the algorithm of Bayati et al. [44] to sample from this distribution.

This experiment allows us to search for structure in the preferential attachment model that is *not* prescribed by the degree distribution. The discrepancy in effectiveness of the normalized versus non-normalized Laplacian distances in Section suggests that much of the structural information that the Laplacian distance is using to discern the two models is contained in the degree distribution. None of the metrics have scaled distances well-separated from zero, which suggests that all significant structural features of the preferential attachment model are prescribed by the degree distribution.

### Watts-Strogatz

In this section, we compare a Watts-Strogatz random graph and an uncorrelated random graph. The Wattz-Strogatz model is interesting in that it contains primarily local structure, in the form of a high local clustering coefficient (i.e. density of triangles).

The Watts-Strogatz model is sparse, and so our volume-matched null model has a low value of *p* and thus is very likely disconnected. This is only a significant problem for the resistance distance, which is undefined for disconnected graphs. To remedy this, we use an extension of the resistance distance called the **renormalized resistance distance**, which is developed and analyzed in [29]. This is the only experiment in which the use of this particular variant of the resistance distance is required.

In Figure 5 we see that the adjacency and normalized Laplacian spectral distances are the strongest performers. Amongst the matrix distanced, DeltaCon strongly outperforms the resistance distance. The resistance distance here shows a negative median, which indicates smaller distances between populations than within the null population. This is likely due to the existence of many (randomly partitioned) disconnected components within this particular null model, which inflates the distances generated by the renormalized resistance distance. It is notable that, contrary to the comparison in Section, the normalized Laplacian outperforms the non-normalized version of the same.

In Figure 7 we see the results for *λ*_*k*_ distances, for a wide variety of *k*. These results indicate that much of the information that the *λ* distances are using to discern between the two models is contained in the higher eigenvalues, particularly for the adjacency and normalized Laplacian distances.

### Lattice Graph

For our final experiment we compare a lattice graph to a random degree-distribution graph with the same degree distribution. The lattice here is highly structured, while the random degree-distribution graph is quite similar to an uncorrelated random graph; both the deterministic degree distribution of the lattice and the binomial distribution of the uncorrelated random graph are highly concentrated around their mean.

We see that the scaled distances in this experiment are about an order of magnitude higher than they are in other experiments for some of the distances; because the lattice is such an extreme example of regularity, it is quite easy for many of the distances to discern between these two models. The resistance distance has the highest performance, while spectral distances all perform equally well. Note that for a regular graph, the eigenvalues of ***A***, ***L***, and ℒ are all equivalent, up to an overall scaling and shift, so we would expect near-identical performance for graphs that are nearly regular.

Similarly to the Watts-Strogatz comparison in Section, much of the information that the *λ* distances use to discern between the models is contained in the higher eigenvalues. This points to the importance of local structure in the lattice.

## Discussion

In this discussion, as we have done throughout the paper, we will emphasize a distinction between local and global graph structure. Global structures include community separation as seen in the stochastic blockmodel, while local structures include the high density of triangles in the Watts-Strogatz model.

In general, we find that when examining global structure, the adjacency spectral distance and DeltaCon distance both provide good performance. When examining community structure in particular, one need not employ the full spectrum when using a spectral distance. The fact that the spectra of the graph provide a natural partitioning [15] aligns with our result that the first few eigenvalues will provide sufficient differentiation if the number of communities is low.

When one is interested in both global and local structure, we recommend use of the adjacency spectral distance. When the full spectrum is employed, the adjacency spectral distance is effective at differentiating between models even if the primary structural differences occur on the local level (e.g. the Watts-Strogatz graph). The use of the entire spectrum here is essential; much of the most important information is contained in the tail of the distribution, and the utility of the adjacency spectral distance decreases significantly when only the dominant eigenvalues are compared.

It is important to remember that these experiments represent only one way that pairwise graph comparison might be used. In particular, we are here comparing a sample to a known population. Alternatively, one might also be interested in comparing a dynamic graph at adjacent time steps [46]; this scenario is treated empirically in Section.

### Discerning Global Structure

Across our models, we see two significant and quite distinct types of global structure, which can be seen in Figure 9. The first of these is the grouping of the graph into distinct communities. The stochastic blockmodel is of course the model which most clearly possesses this type of global structure. At the local level, the stochastic blockmodel is nearly identical to the uncorrelated random graph, and so we can use the results of Section to understand how distances respond to this specific feature.

The particular configuration of the stochastic blockmodel that we use has two partitions of equal size. We would thus expect the second eigenvalue *λ*_2_ to be the primary distinguishing spectral feature of the graph (in any of the three matrix representations used). Indeed we see in Figure 1 that this is the case, and that the use of additional eigenvalues beyond *k* = 2 only serves to decrease performance by including noise in the comparison. In Figure 10 we can directly observe the similarity in spectra between the two models, as well as the presence in the stochastic blockmodel of a second eigenvalue which separates from the bulk of the spectrum.

The separation of the first *k* eigenvalues from the semi-circular bulk spectrum of the stochastic blockmodel is studied analytically in [43]. The authors show that a graph spectrum can be though of as two distinct components; a continuous bulk, and discrete outliers, with the latter indicating community structure. This separation is what allows our *λ*_*k*_ distances to function effectively in detecting community structure. In general, the use of the spectrum for community partitioning in graphs has a long history [47]. Recent, Lee et al. [15] have proven a performance bound on the effectiveness of using the first *k* eigenvectors to partition the graph into *k* clusters.

In [29], the authors study the performance of the resistance perturbation metric in the setting of a dynamic variant of the stochastic blockmodel. Although not the same scenario as ours, their result is closely connected, and conforms well with the currently observed data. In particular, their result indicates that for the resistance metric to be effective in detecting topological changes in a stochastic blockmodel, the number of cross-community edges must be asymptotically dominated by the mean degree.

This is a highly restrictive condition. In the results shown in Section, we see that the resistance metrics performs poorly; auxiliary results (not shown) indicate that its performance increases significantly when the graph has high in-community degree *p* = 0.495 and very low cross-community connection *q* = 0.005. This unrealistic density requirement puts severe practical restrictions on the applicability of using the resistance metric to detect topological changes in community graphs. Furthermore, in these extreme cases, other measures (such as the spectral distances) can also easily distinguish between the two models.

The link between graph resistance and degree has been established in [48], where the authors show that the resistance *R*_*uv*_ between vertices *u* and *v* can be well approximated by

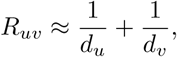

an approximation which suggests that fluctuations in degree distribution would result in significant fluctuations in the graph resistance. This is corroborated by the poor performance of the resistance distance in Section. These results indicate that the resistance distance cannot “see” changes in global structure over local noise, unless the global structure is unrealistically stark (as in the asymptotic condition given in [29]).

The second significant global structure seen in our models is the particular topology in which there are a small number of highly connected vertices which dominate the connectivity patterns of the graph. A small graph exhibiting this structure can be seen in Figure 9. This results in a heavy-tailed degree distribution. The random graph model which features this type of structure is the preferential attachment model, whose degree distribution exhibits polynomial decay in the tails [9].

The best tool for detecting this structure is the Laplacian spectral distance. The presence of the degree matrix *D* in the Laplacian *L* = *D – A* means that comparison of Laplacians is very effective for discerning between models with radically different degree distributions. Since significant differences between the degree distributions of the preferential and attachment graphs occur in the tail (i.e. high-degree vertices), the inclusion of the final few eigenvalues is essential if one wishes to use the Laplacian spectrum to perform this comparison.

Figure 10 exhibits the influence of the degree distribution on the Laplacian spectrum. We observe qualitatively, as demonstrated in [19], that the tails of the Laplacian spectrum of a preferential attachment graph exhibits polynomial decay similar to the tail of the degree distribution. This is a prime example of the way in which the spectrum of the Laplacian can be heavily influenced by the degree distribution.

The particular topology of the preferential attachment differentiates itself from that of the uncorrelated random graph at both a global and local level. Even though the Laplacian spectral distance is best at observing the significant effect of high-degree vertices on the model, it is not, all in all, the most efficient tool for differentiating the two. To understand this, let us now turn to further discussion of the local structure present in the preferential attachment model, as well as the other models studied.

### Impact of Local Structure

Local structure consists of structures existing at the level of a single vertex or subgraphs consisting of a small number of vertices. These local structures can provide important information about the topology of the graph, or they can amount to noise which obfuscates our ability to examine global structures of interest. Our experiments provide examples of both of these cases, which we will now examine.

Consider first the results of Section. In Figure 4, we see that the adjacency spectral distance differentiates between the two models based primarily on the first dominant eigenvalue. Recalling the interpretation of the adjacency spectrum provided by Maas [24] and reiterated in Section, we realize that this is due to the high density of low-degree vertices in the preferential attachment model, compared to the uncorrelated random graph. This local structure is in some sense necessitated by the presence of a few very high degree vertices, since we demand that the graphs being compared are equal in expected volume. Indeed, the degree distribution of this model is so structurally significant that it almost entirely determines the structure of the resulting graph. We see in Figure 3 that no distances can effective discern between a preferential attachment graph and a randomized graph with the same degree distribution.

The Watts-Strogatz graph is another example of a model whose signature lies primarily in local structure. Farkas et al. [19] argue that the presence of a high number of triangles is the distinguishing feature of a Watts-Strogatz graph, and persists at values of *β* in which other structural aspects of the ring lattice (e.g. regularity and periodicity). The third moment of the spectral density of ***A*** tells us the expected number of triangles in a graph,^12^ and so one would expect inclusion of the full spectrum important in detecting the topological signature of this model. On a global scale, the model does not significantly differ from the uncorrelated random graph; highly connected vertices are extremely unlikely, and the generative rewiring mechanism does not result in the presence of communities in the graph.

We see in Figure 7 that inclusion of the large-*k* (high frequency) eigenvalues is essential to differentiating between the models. In Figure 10 we see that the spectral density of the Watts-Strogatz model exhibits high skewness, which indicates the high expected number of triangles in the graph, and is only captured by inclusion of the full spectral bulk.

The lattice graph is an extreme example of this kind of local structure. Similarly to the Watts-Strogatz model, there is a ubiquity of a certain type of local structure in the graph, namely the presence of four-edge loops. We see in Figure 8 that including a large number of eigenvalues in spectral comparison greatly increases the efficacy of the spectral distances. However, since the lattice is so remarkably regular (unlike the Watts-Strogatz model, which is a perturbed ring lattice) even comparing only a few principal eigenvalue is sufficient to differentiate it from a randomized graph with the same degree distribution. The spectra of the two models are shown in Figure 10.

Local structure is sometimes important when understanding graph structure, but can also frequently serve as a source of uninformative noise when comparing graphs. The results of Section illustrate this fact. Looking to Figure 1, we see that the Laplacian spectrum is unable to distinguish between the stochastic blockmodel and the uncorrelated random graph, while the normalized Laplacian distinguishes them well. The difference between these two matrix representations is that normalization removes degree information, which is not informative in this particular model.

We see a similar problem arise when we apply the resistance distance to the stochastic blockmodel; as discussed in the previous section, the resistance distance is disproportionately influenced by local structure, and is unable to discern the global structure of the graph over local fluctuations. Interestingly, DeltaCon does not appear to suffer from local fluctuations as much as the resistance distance. This could be due to the structure of the matrix **S** that DeltaCon uses to represent the graph, or due to the use of the Matusita distance rather than the *ℓ*_1_ or *ℓ*_2_ norm to compare the resulting matrices (for more discussion of this, see Sections 2.2 and 3.1 in [5]).

It is essential to determine whether local topological features are of interest in the comparison problem at hand; inclusion of locally targeted distance measures can hinder the performance of graph distances in cases where local structure is noisy and uninformative. However, if local structure is ignored, one can often omit essential structural information about the graphs under comparison.

### Recommendations

Throughout our experiments, the most consistent observation is that the adjacency spectral distance shows high effectiveness in discerning between a variety of models. We see in Section that it is able to perceive global community structure and not be overwhelmed by local fluctuations in degree, but Sections and show that it is by no means ignorant of local structure present in a graph. That is to say, the adjacency spectral distance is *multiscale*; the scale of interest can be chosen by tuning the number of principal eigenvalues included in the comparison.

Spectral distances exhibits practical advantages over matrix distances, as they can inherently compare graphs of different sizes and can compare graphs without known vertex correspondence. The adjacency spectrum in particular is well-understood, and is perhaps the most frequently studied graph spectrum; see e.g. [19, 41]. Finally, fast, stable eigensolvers for symmetric matrices are ubiquitous in modern computing packages such as ARPACK, NumPy, and Matlab, allowing for rapid deployment of models based on spectral graph comparison.^13^ Furthermore, randomized algorithms for matrix decomposition allow for highly parallelizable calculation of the spectra of large graphs [34].

However, the utility of the adjacency spectral distance is not general enough to simply apply it to any given graph matching or anomaly detection problem in a naive manner. A prudent practitioner would combine exploratory structural analysis of the graphs in question with an ensemble approach in which multiple distance measures are considered simultaneously, and the resulting information is combined to form a consensus. Such systems are commonplace in problems of classification in machine learning, where they are sometimes known as “voting classifiers” (see e.g. [49]).

As we have said before, we have been comparing graphs of equal volume (in expectation). In situations where the graph volume varies drastically, the process of choosing a graph comparison tool may differ significantly. We will address this in Section, where we deal with graphs that exhibit significant volume fluctuations.

## Applications to Empirical Data

Random graph models are often designed to simulate a single important feature of empirical networks, such as clustering in the Watts-Strogatz model or the high-degree vertices of the preferential attachment model. In empirical graphs, these factors coexist in an often unpredictable configuration, along with significant amounts of noise. Although the above analysis of the efficacy of various distances on random graph scenarios can help inform and guide our intuition, to truly understand their utility we must also look at how they perform when applied to empirical graph data.

In this section, we will examine the performance of our distance in two scenarios. First, we will look at an anomaly detection scenario for a dynamic social-contact graph, collected via RFID tags in an French primary school [50]. Secondly, we will look at a graph matching problem in neuroscience, comparing correlation graphs of brain activity in subjects with and without autism spectrum disorder [51].

The first experiment suggests that the tools that perform the most consistently in the graph matching applications (the spectral distances) are unreliable in our anomaly detection experiment. It is also interesting insofar as the graphs exhibit significant volume fluctuations, which was a factor not present in our numerical studies.

In the second experiment, we see that none of our graph distances fully distinguish between the two populations. Signal-to-noise is a ubiquitous problem in analyzing actual graph data, and is particularly notable in building a connectivity networks of human brain activity (see e.g. [52]). Accordingly, the results of our data experiments show that in the presence of real-world noise levels, many of these distances fail to distinguish subtle structural differences. In the face of this, we examine more targeted analysis techniques which may be applied in such a situation.

### Primary School Social Contact Data

Some of the most well-known empirical network datasets reflect social connective structure between individuals, often in online social network platforms such as Facebook and Twitter. These networks exhibit structural features such as communities and highly connected vertices, and can undergo significant structural changes as they evolve in time. Examples of such structural changes include the merging of communities, or the emergence of a single user as a connective hub between disparate regions of the graph.

In this section, we investigate a social contact network, which is based on measurements of face-to-face contact using RFID tags. We use our distances to compare the graph at subsequent time steps. This is a quite different scenario than that presented in Section; the most important difference is that there is a natural sense of vertex correspondence, because the students’ labels persist over time. This change has significant implications for the performance of our various distances, which we will explore in the discussion below.

#### Description of Experiment

The data are part of a study of face to face contact between primary school students [50]. Briefly, RFID tags were used to record face-to-face contact between students in a primary school in Lyon, France in October, 2009. Events punctuate the school day of the children (see Table 3), and lead to fundamental topological changes in the contact network (see Fig. 11). The school is composed of ten classes: each of the five grades (1 to 5) is divided into two classes (see Fig. 11).

**Table 3.** Events that punctuate the school day.

| Time | Event |
| --- | --- |
| 10:30 a.m. – 11:00 a.m. | Morning Recess |
| 12:00 p.m. – 1:00 p.m. | First Lunch Period |
| 1:00 p.m. – 2:00 p.m. | Second Lunch Period |
| 3:30 p.m. – 4:00 p.m. | Afternoon Recess |

**Fig 11.**
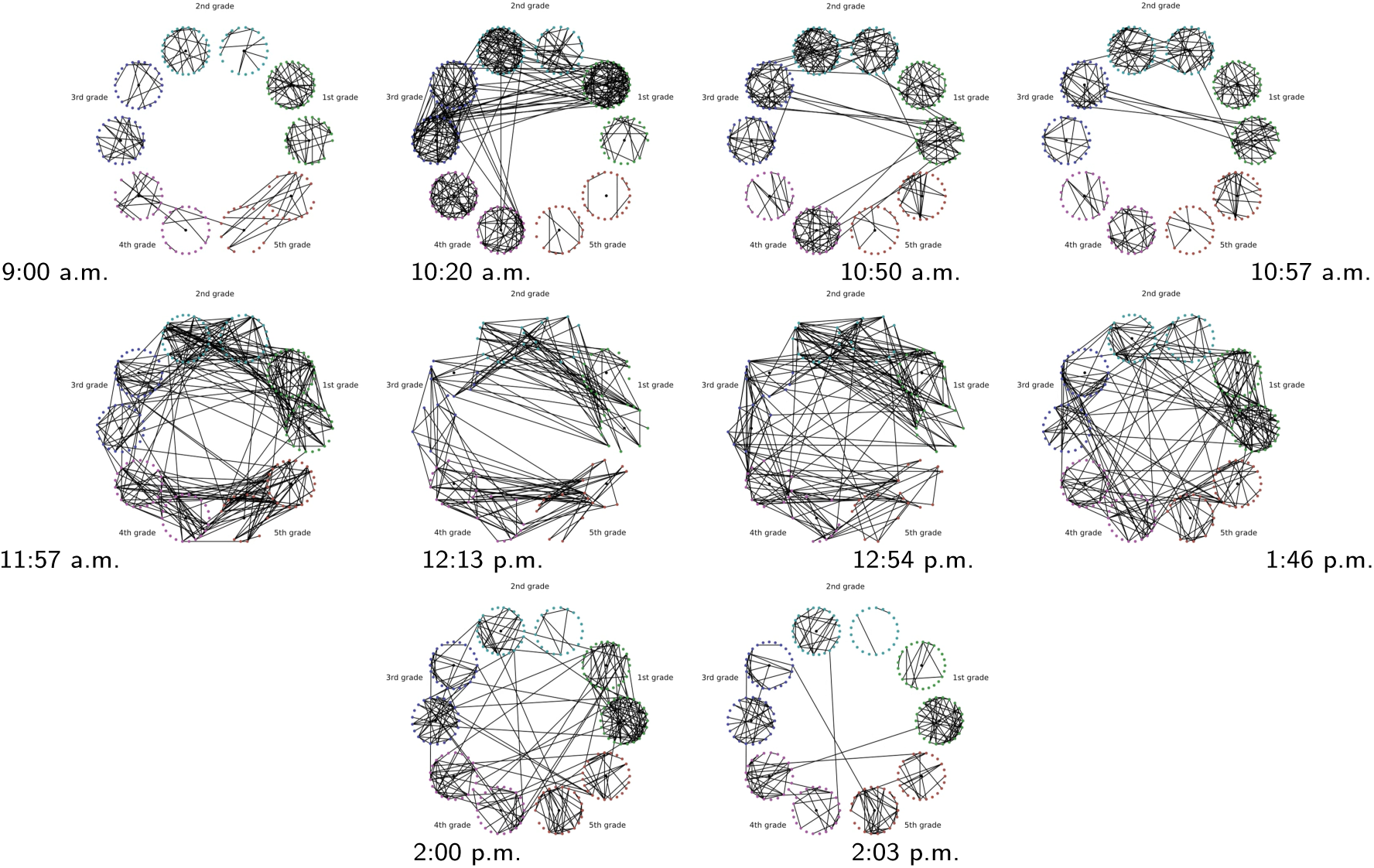
Top to bottom, left to right: snapshots of the face-to-face contact network at times (shown below each graph) surrounding significant topological changes.

The construction of a dynamic graph proceeds as follows: time series of edges that correspond to face to face contact describe the dynamics of the pairwise interactions between students. We divide the school day into *N* = 150 time intervals of Δ*t ≈* 200 s. We denote by *t*_*i*_ = 0, Δ*t, …*, (*N -* 1)Δ*t*, the corresponding temporal grid. For each *t*_*i*_ we construct an undirected unweighted graph 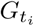, where the *n* = 232 nodes correspond to the 232 students in the 10 classes, and an edge is present between two students *u* and *v* if they were in contact (according to the RFID tags) during the time interval [*t*_*i-*1_, *t*_*i*_).

Changes in the graph topology during the school day are quantified using the various distance measures,

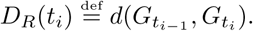

To help compare these distances with one another, we normalize each by their sample mean 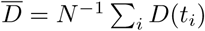, and we define

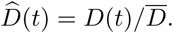

For the purpose of this work, we think of each class as a community of connected students; classes are weakly connected (e.g., see Fig. 11 at times 9:00 a.m., and 2:03 p.m.). During the school day, events such as lunchtime and recess, trigger significant increases in the the number of links between the communities, and disrupt the community structure; see Fig. 11 at times 11:57 a.m., and 1:46 p.m..

## Discussion

In Figure 12, we see the normalized time series 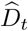 for each of the distance measures studied. Interestingly, the matrix distances all achieve passable performance, while NetSimile and the spectral distances are far too noisy to be of any practical use. As we see in Figure 11, the main structural changes that the graph undergoes are transitions into and out of a strong ten-community structure that reflects the classrooms of the school. For example, the adjacency matrix begins as (mostly) block-diagonal at 9 AM, but has significant off-diagonal elements by morning recess at 10:20 AM, and is no longer (block) diagonally dominant come the lunch period at 12 PM.

These structural changes are of a global nature. In Section we saw that the spectral distances were more effective than the matrix distances at detecting differences in community structure between graphs. Why is this not the case here? The graphs are persistent, in the sense that the vertices show a natural correspondence, which the matrix distances exploit. For example, we know that certain edges (those between classes) are “cross-community,” and so the presence of these edges suggests some anomalous topology. This is why even the edit distance is quite effective at detecting topological changes in the graph.

Now let us highlight certain interesting features of the comparison shown in the top plot in Figure 12. Amongst the matrix distances, the resistance distance shows the largest distance at 10:20 AM, followed by the smallest distance at the subsequent time step. During recess, local changes are occurring in the graph, but the global structure is remaining mostly constant – the graph consists of strongly connected communities, with some connection between them. The fact that the resistance distance shows the lowest distance between time steps within recess suggests that, of the matrix distances, it is least effected by this local variation in the graph.

The lunch periods are marked by a stark change in graph topology. The graph undergoes significant global structural transformation, becoming almost entirely unordered with respect to class communities. Again the resistance distance shows a more significant signal at the beginning of this change, and a smaller signal as the anomalous topology persists.

The next significant transformation the graph undergoes is the transition out of lunch periods. The resistance distance stands out as marking this transition most pronouncedly, although all distances show significantly smaller distances between the time steps during the afternoon class period compared to during the lunch period.

Unlike our numerical experiments above, the graphs being compared here show significant fluctuations in volume. However, these fluctuations do not align with our event markers to the extent that our matrix distances do. This indicates that our matrix distances are picking up more than simply changes in volume.

The most remarkable conclusion of this particular experiment is that although the spectral distances are very efficient and stable for the purposes of graph matching, they show very poor performance in anomaly detection on dynamic graphs. This is due to the inherent vertex correspondence that is automatically provided when comparing a dynamic graph at subsequent time steps. Although we reflect on the subtle distinctions in performance between the three matrix distances, they all show very similar overall performance, and any one of them would be sufficient for application in this scenario.

### Brain Connectomics of Autism Spectrum Disorder

Graph theoretical analysis of the connective structure of the human brain is a popular research topic, and has benefited from our growing ability to analyze network topology [1, 53]. In these graph representations of the brain, the vertices are physical regions of the brain, and the edges indicate the connectivity between two regions. The connective structure of the brain is examined either at the “structural” level, in which edges represent anatomical connection between two regions, or at the “functional” level, in which an edge connects regions whose activation patterns are in some sense similar.

Psychological conditions such as Alzheimer’s disease [54], autism spectrum disorder [55], and schizophrenia [56] have been shown to have structural correlates in the graph representations of the brains of those affected. In this section, we will focus on autism spectrum disorder, or ASD. The availability of high-quality, open-access preprocessed data [57] makes ASD a particularly attractive choice for researchers with little experience implementing the nuanced preprocessing pipelines seen in neuroimaging. We examine which, if any, of our graph distances are able to effectively discern between subjects with ASD and subjects which are typically developing (TD). This is a problem in graph matching, very similar in structure to the experiments done on random graph models above.

We will see that our distances are ineffective at discerning between the graphs arising from ASD subjects and TD subjects. This result agrees with the conclusion of a recent review, which finds that classification methods do not generalize well to novel data [58]. The negative outcome of this experiment both informs an understanding of the limitations of generalized tools such as our graph distances, and points to possible refinements of these tools.

#### Description of the Data

The Autism Brain Imagine Data Exachange, or ABIDE, is an aggregation of brain-imaging data sets from laboratories around the world which study the neurophisiology of ASD [51]. The data that we focus on are measurements of the activity level in various regions of the brain, measured via functional magnetic resonance imaging (fMRI). The fMRI method is actually measuring blood oxygen levels in the brain, which are then used as a proxy for activation levels. Measurements are taken over myriad small volumes within the brain, preprocessed, and then aggregated into a much smaller collection of time series, each representing a distinct region of the brain.

These time series then pass through an extensive preprocessing pipeline, which includes myriad steps such as nuisance signal (e.g. heartbeat) removal, detrending, smoothing via band-pass filtration, and so on. A detailed assessment of the preprocessing steps can be found in [57]. After preprocessing, the data is analyzed for quality. Of the original 1114 subjects (521 ASD and 593 TD), only 871 pass this quality-assurance step. These subjects are then spatially aggregated via the Automated Anatomical Labelling (AAL) atlas, which aggregates the spatial data into 116 time series.

To construct a graph from these time series, the pairwise Pearson correlation is calculated to measure similarity. If we let *u* and *v* denote two regions in the AAL atlas and let *P* (*u, v*) denote the Pearson correlation between the corresponding time series, the simplest way to build a graph is to assign weights *w*(*u, v*) = |*P* (*u, v*) |, so the weight between vertices *u* and *v* is the modulus of the correlation. One may wish to exclude particularly low correlations, as these are often spurious and not informative as to the structure of the underlying network. In this case, one chooses a threshold *T* and assigns weights via

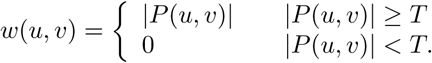

Finally, one may wish to binarize the graph, so that *w ∈ {*0, 1}, and thus uses the formula

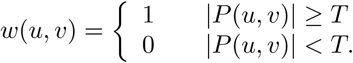

We will compare both binary and weighted connectomes, generated at multiple thresholding levels. This will allow us to be confident that our results are not artifacts of poorly chosen parameters in our definition of the connectome graph.

## Discussion

Figure 13 shows the results of our experiment comparing connectomes of TD and ASD subjects. The TD subjects play the role of the null population ***𝒢***_1_ and the ASD subjects constitute the alternative population ***𝒢***_2_, and the scaled distances shown in the plot are calculated in the manner outlined in Section.

**Fig 12.**
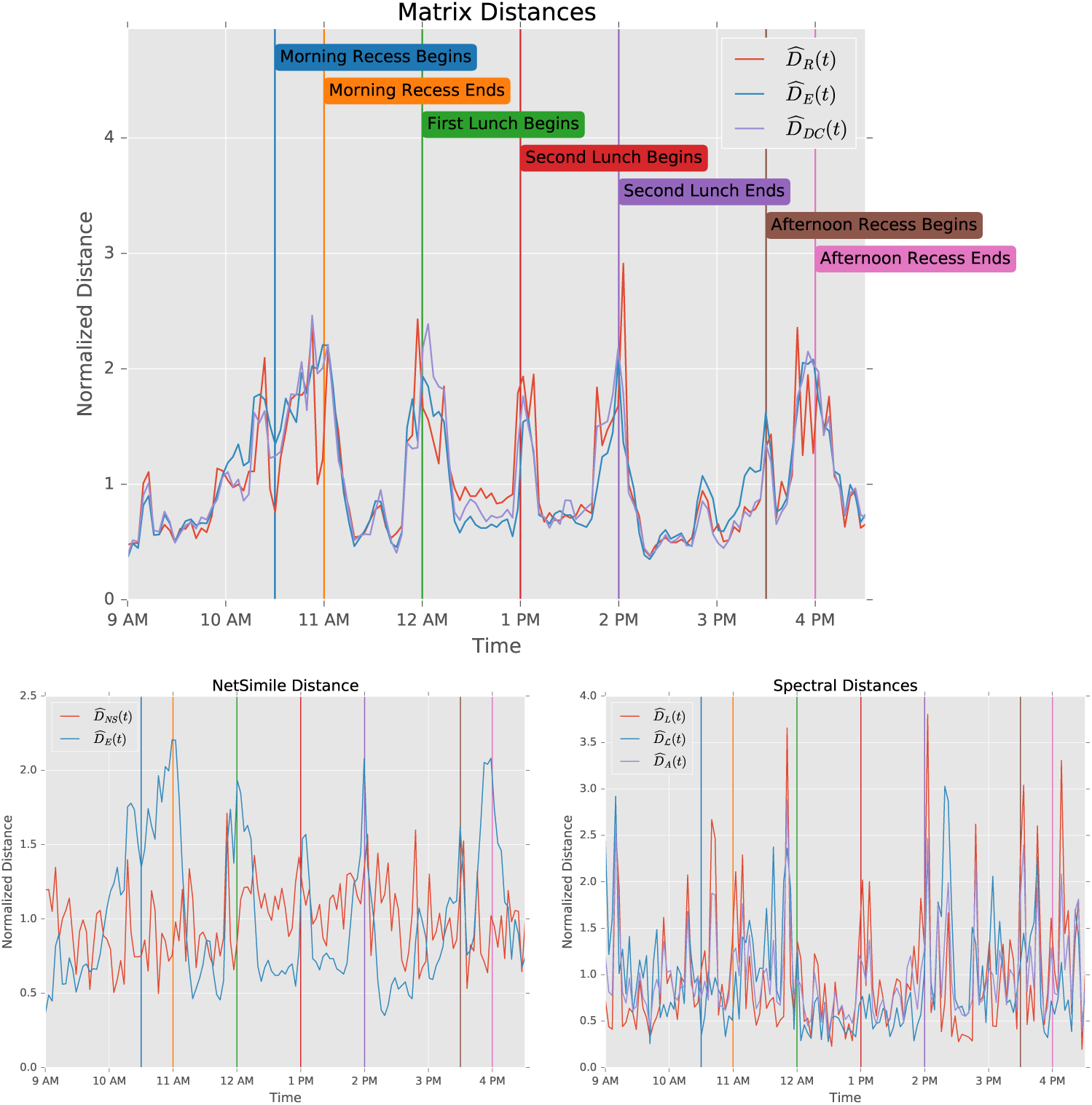
Comparison of distance performance on primary school data set. All distances are normalized by their sample mean.

**Fig 13.**
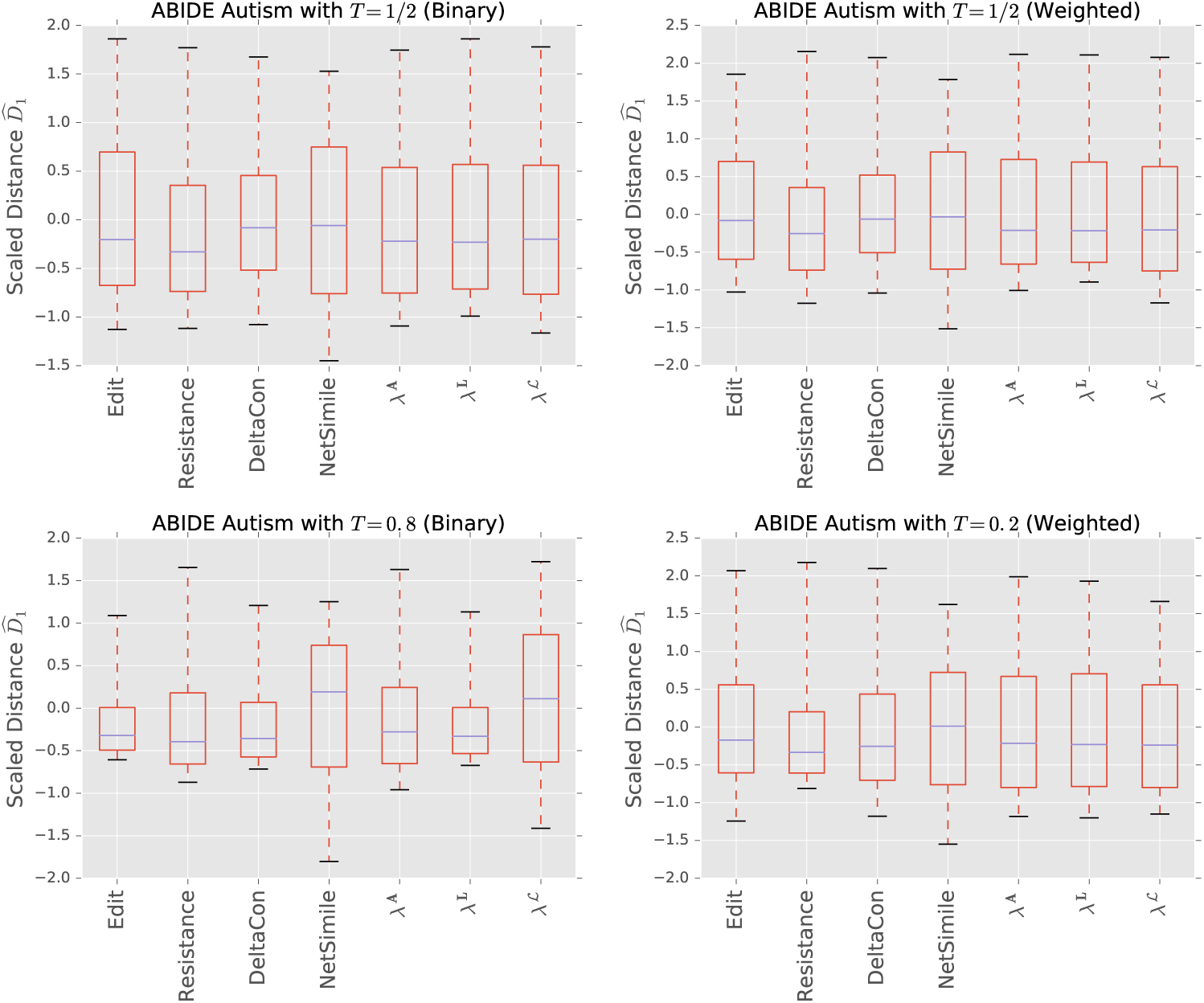
Distance comparison for ABIDE autism data set, for various thresholding configurations.

We observe that no distances effectively separate the two communities, regardless of level of thresholding, or the presence or absence of binarization. Indeed, the negative median of the scaled distance indicates that distances from ASD to TD connectomes is *lower* than distances between two TD connectomes, indicating a higher structural variability in TD connectomes. As we will see below, the structural differences between the two communities are localized within subgraphs, and do not persist throughout the full graph. Furthermore, the signal produced by these differences is not easily differentiated from the local variations (i.e. noise) present in the communities. For these two reasons, global comparison using graph metrics is ineffective for this problem.

Figure 14 shows a region-by-region comparison of connectomes of ASD and TD subjects. Similarly to our previous scaling, we take the mean and variance (region-wise) of the correlations in TD subject, and then use these to normalize the correlations of ASD subject. Thus, a value of 0.25 in Figure 14 indicates that ASD subject show, on average, correlations 0.25 standard deviations above the mean, relative to TD subjects.

**Fig 14.**
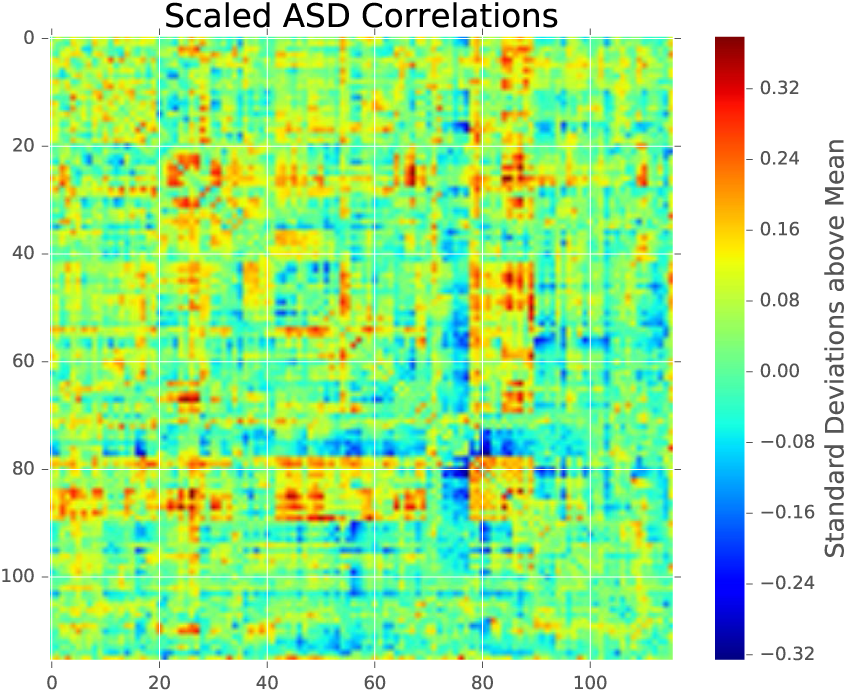
Absolute correlations between regions in ASD subjects. Data are normalized by mean and variance of absolute correlations between TD subjects.

We see that certain regions show significant differences. A closer examination shows that ASD subjects are generally underconnected in regions 73 through 77, and are overconnected in regions 79 and 84 through 89. A table below shows the specific anatomical regions that these labels correspond to.

**Table 4.**
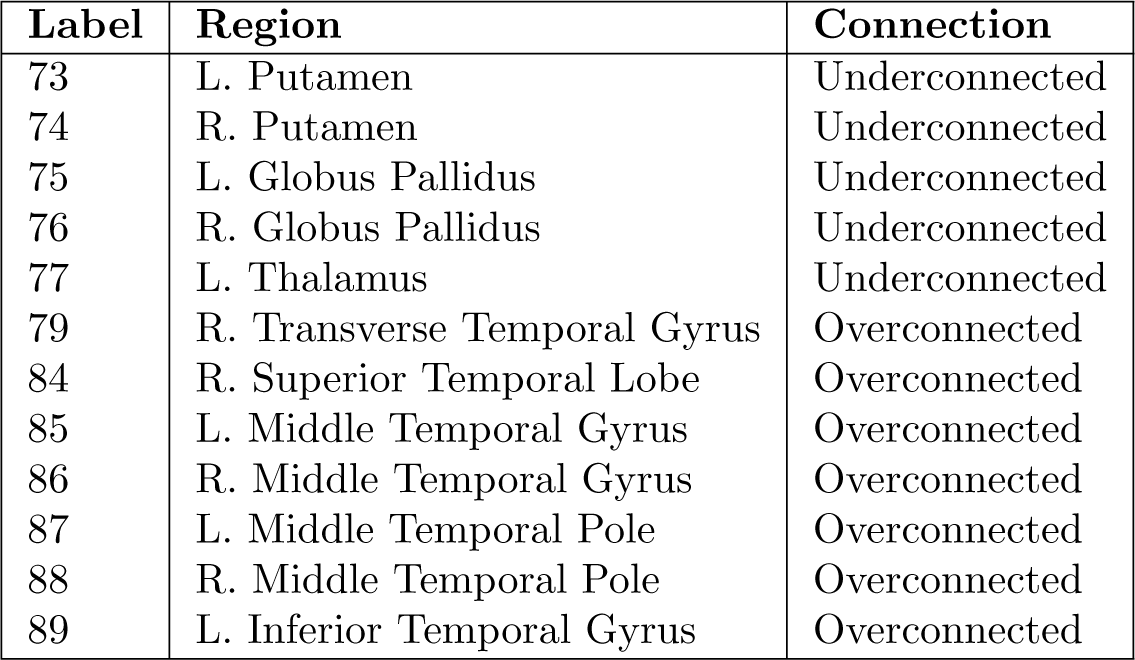
Regions that show notably anomalous connectivity patterns. Correspondence between labels and regions is established via the Automated Anatomical Labelling atlas [59].

| Label | Region | Connection |
| --- | --- | --- |
| 73 | L. Putamen | Underconnected |
| 74 | R. Putamen | Underconnected |
| 75 | L. Globus Pallidus | Underconnected |
| 76 | R. Globus Pallidus | Underconnected |
| 77 | L. Thalamus | Underconnected |
| 79 | R. Transverse Temporal Gyrus | Overconnected |
| 84 | R. Superior Temporal Lobe | Overconnected |
| 85 | L. Middle Temporal Gyrus | Overconnected |
| 86 | R. Middle Temporal Gyrus | Overconnected |
| 87 | L. Middle Temporal Pole | Overconnected |
| 88 | R. Middle Temporal Pole | Overconnected |
| 89 | L. Inferior Temporal Gyrus | Overconnected |

Figure 14 indicates that there are in fact significant structural differences between the connectomes of TD and ASD subjects. However, the differences barely stand out above the noise in the graph; all the edge differences in Figure 14 are less than half a standard deviation away from the mean. Furthermore, the differences occur in isolated regions of the graph, and the majority of the edge weights do not show statistically significant differences between the two populations. Both the low amplitude and small extent of the signal contribute to the difficulty we see graph distances have in discerning between TD and ASD subjects.

In [60], the authors find very little that distinguishes the connectomes of ASD subjects from TD subjects, save for lower betweenness centrality in right later parietal region. Similarly, the authors of [58] assert that although classification algorithms show “modest to conservatively good” accuracy rates, they perform poorly when tested on novel data sets. Looking in particular to [61], we see that the authors are able to achieve just over 75% accuracy in their classification, but they do so by both preprocessing the connectomes via particular regions of interest, and by testing a smorgasbord of classifiers (9 different models are used) to find the one which shows highest performance. Such a process exhibits that reasonable accuracy can be achieved via careful algorithmic tuning; inversely, our result shows that a naïve application of a graph-theoretic method does not provide us with accurate classification.

A complete explanation of the failure of connectomes to unambiguously characterize ASD is well beyond the scope of this work; however, we will highlight a few interesting possibilities. In [61], the authors suggest that poor performance of many classifiers may be due to inclusion of uninformative features. Said another way, regions of the brain that have no bearing on the presence or absence of ASD are included in the connectome, which then reduces the signal to noise ratio of the data. In [58], the authors raise the issue of mental state in which the scans are performed. This state is often referred to as a “resting state,” but variations in instructions (i.e. eyes open vs. closed) can have strong effects on the resulting data [62]. In [63], the authors find that when ASD connectomes show lower connectivity than TD connectomes when the subject exteroceptive activity, but show *higher* connectivity during introspective attentional tasks. This contrast highlights the need for careful control of the subjects’ mental state if meaningful comparisons between connectomes are to be made.

It should also be noted that the choice of Pearson correlation to compare the fMRI time series is not obviously the correct one. It has been recently shown that the time series of brain activity exhibit nontrivial lag structure [64], indicating the need for a more general method of time series comparison. Myriad popular methods such as Granger causality [65] and mutual information can be applied to this problem; indeed, mutual information analysis has already shown promise as a tool in the pre-processing and feature selection stage of connectome analysis [66].

## Conclusion

We have studied the efficacy of various graph distance measures when they are used to differentiate between popular random graph models, as well as in empirical anomaly detection and graph classification scenarios. These measures are understood through a multi-scale lens, in which the impact of global and local structures are considered separately. Although recent work [67] has called into question the previously assumed ubiquity of some of these models, studying their properties builds qualitative and guide intuition when examining empirical network datasets.

Throughout our graph matching experiments in Section, we find that the adjacency spectral distance is the most stable, in the sense that it exhibits good performance across a variety of scenarios. It exhibits an ability to perceive both global and local structure, while avoiding becoming overwhelmed by local fluctuations in graph topology. Although the various matrix distances we examine allow for elegant analysis [5, 11], we find their performance on random graph model comparison underwhelming.

The situation reverses when we look at dynamic anomaly detection in Section. In this scenario, the matrix distances proved most effective, and showed clear indications of the ground-truth anomalies present in the data. The spectral distances, on the other hand, were so noisy as to be useless. When doing anomaly detection on a dynamic graph, the two graphs in comparison tend to share more edges than in graph matching scenarios, which may contribute to the good performance of our matrix distances. Although the graphs in this section fluctuate in volume, we do not find that these fluctuations are helpful in detecting anomalous time steps.

Finally, we explore a collection of human connectomes of subjects with and without autism spectrum disorder. We observe that although differences are observable within the two populations via statistical comparison of edge weights, no graph distance effectively separates the two populations. This experiment helps us understand the limitations of using such generalized tools. We conclude that either more targeted tools are necessary, or more careful data collection and preprocessing is needed to establish a dataset that is separable via classification algorithms.

Based on the results of our numerical and empirical data experiments, we provide a suggested decision process in Figure 15. If the graphs to be compared exhibit differences in volume or size, then these should be examined to see if they hold predictive power, as they are so simple and efficient to compute. If they prove ineffective, then one must consider the setting. In a dynamic setting, in which a dynamic graph is being compared at subsequent time steps, then we recommend using matrix distances based on the results of Section. If one is comparing graphs to determine whether a sample belongs to a given population, then the adjacency spectral distance is the most reliable, as Section demonstrates. Finally, if none of these approaches give adequate performance, then a more targeted analysis must be performed, such as the edge-wise statistical comparison of weights in Figure 14. The particular design of this analysis is domain specific and highly dependent upon the nature of the data.

**Fig 15.**
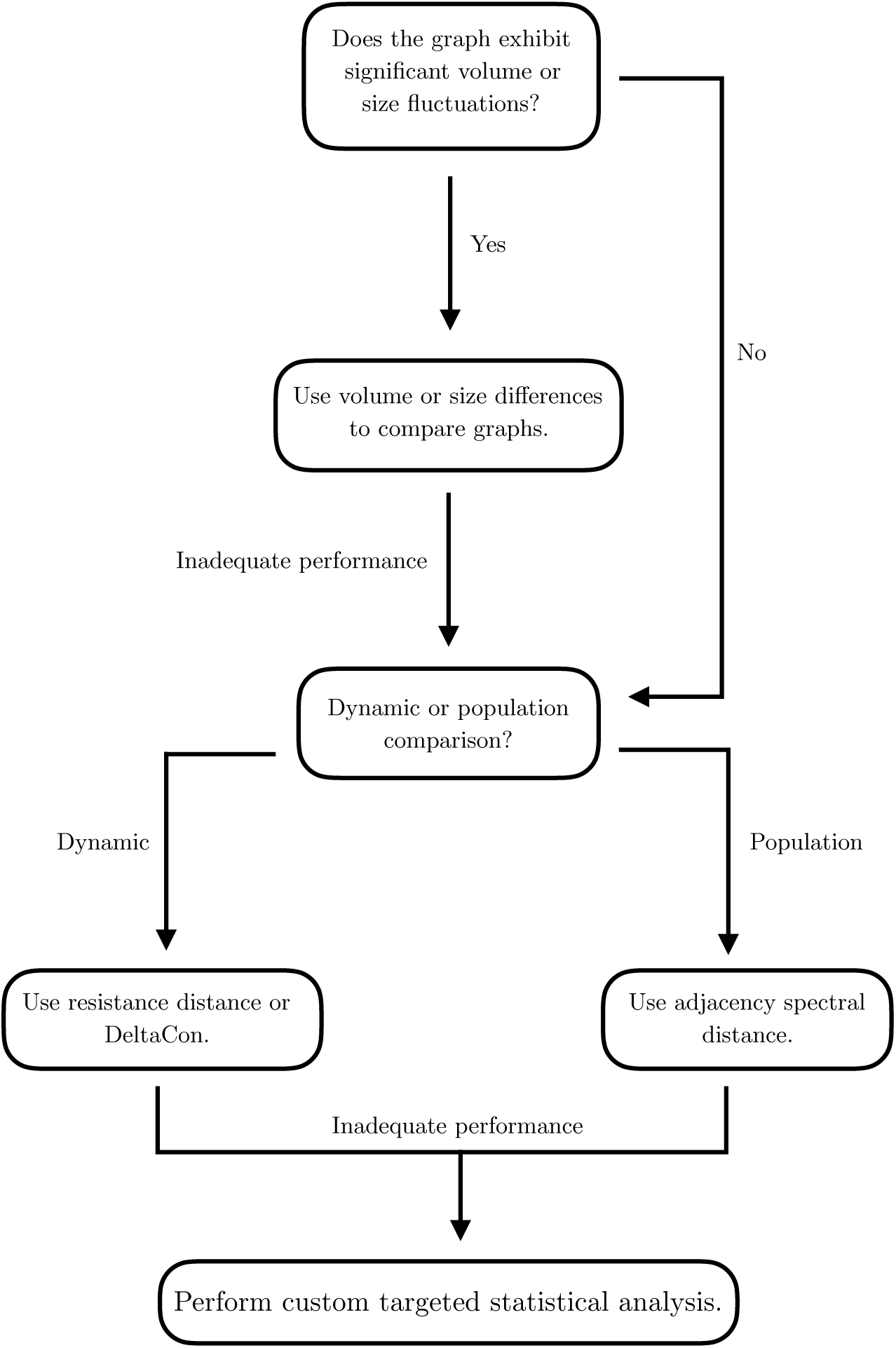
Flow chart summarizing the suggested decision process for applying distance measures in empirical data.

## Notation

For reference, in Table 5 we provide a table of notation used throughout the paper.

**Table 5.**
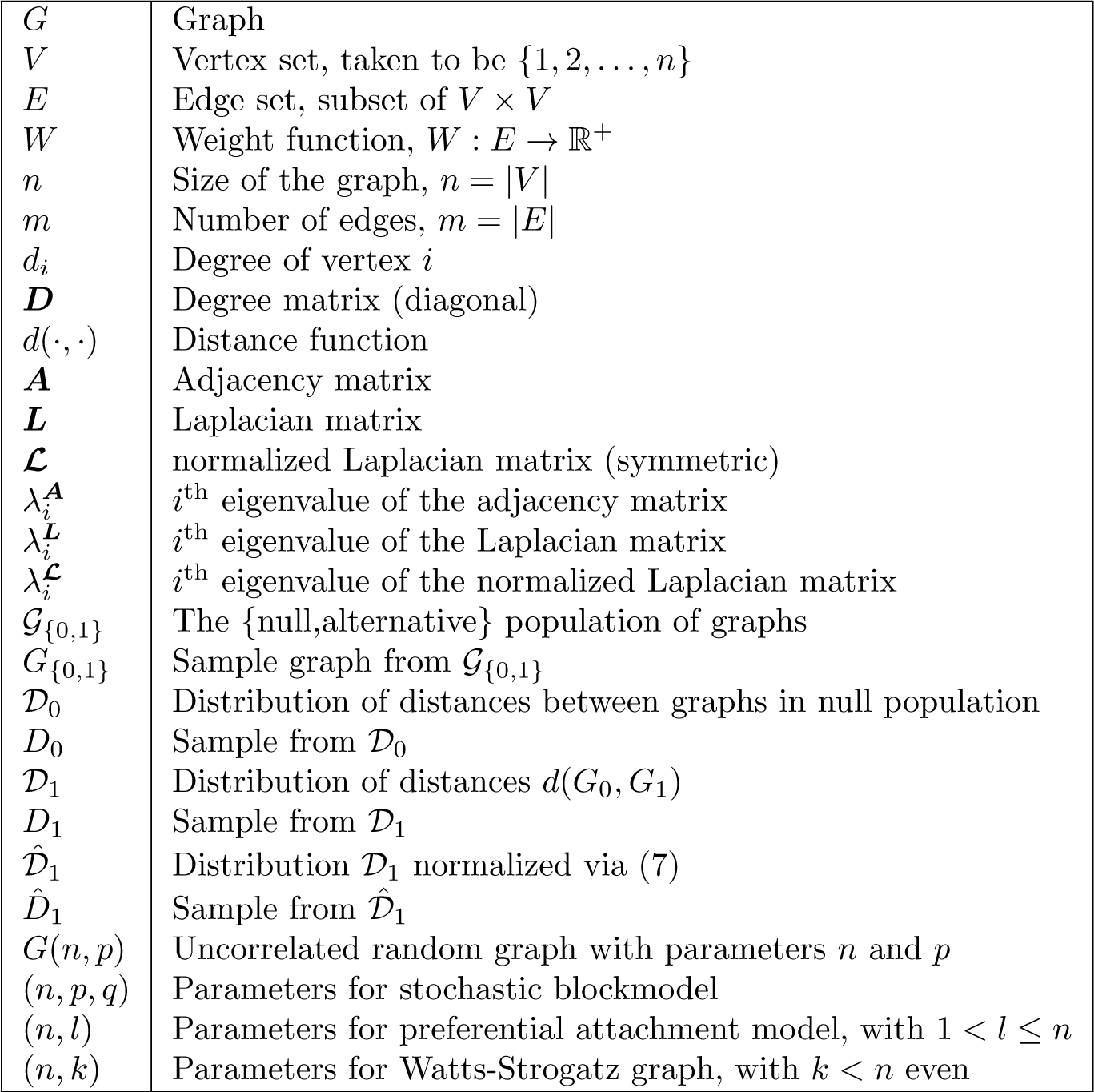
Table of commonly used notation.

## NetComp: Network Comparison in Python

`NetComp` is a Python library which implements the graph distances studied in this work. Although many useful tools for network construction and analysis are available in the well-known `NetworkX` [38], advanced algorithms such as spectral comparisons and DeltaCon are not present. `NetComp` is designed to bridge this gap.

### Design Consideration

The guiding principles behind the library are

#### 1. Speed

The library implements algorithms that run in linear or near-linear time, and are thus applicable to large graph data problems.^14^

#### 2. Flexibility

The library uses as its fundamental object the adjacency matrix. This matrix can be represented in either a dense (`NumPy` matrix) or sparse (`SciPy` sparse matrix) format. Using such a ubiquitous format as fundamental allows easy input of graph data from a wide variety of sources.

#### 3. Extensibility

The library is written so as to be easily extended by anyone wishing to do so. The included graph distances will hopefully be only the beginning of a full library of efficient modern graph comparison tools that will be implemented within `NetComp`.

`NetComp` is available via the Python Package Index, which is most frequently accessed via the command-line tool `pip`. The user can install it locally via the shell command

`pip install netcomp.`

As of writing, the library is in alpha. The approximate (near-linear) forms of DeltaCon and the resistance distance are not yet included in the package. Both algorithms have an quadratic-time exact form which is implemented. Those interested can view the source code and contribute at https://www.github.com/peterewills/netcomp.

## Acknowledgments

F.G.M was supported by the National Natural Science Foundation (CCF/CIF 1815971), and by a Jean d’Alembert Fellowship.

## Author contributions

Writing – Original Draft: P.W.; Writing – Review & Editing: P.W. and F.G.M.; Conceptualization: P.W. and F.G.M.; Investigation: P.W. and F.G.M.; Methodology: P.W. and F.G.M; Formal Analysis: P.W. and F.G.M.; Software: P.W.

## Footnotes

1 These objects are sometimes referred to as “complex networks;” we will use the mathematician’s term “graph” throughout the paper.

2 Sparsity is, roughly, the requirement that the number of edges in a graph of size *n* be much lower than the maximum possible number *n*^2^/2; a technical definition is provided below.

3 Note that the authors in [8] are classifying anomaly detection methods in particular, rather than graph comparison methods in general.

4 “Spectral structure” might refer to the overall shape of the spectral density, or the value of individual eigenvalues separated from the bulk.

5 If the graph is already regular with degree *d*, then this interpretation is consistent with the previous, since the eigenvalues of ***L*** = *d****I*** – ***A*** are just 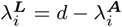.

6 When we say “distance” we implicitly assume that smaller values imply greater similarity; however, we can also carry out this approach with a similarity score, in which larger values imply greater similarity.

7 We could use metrics, or even similarity functions here, although that may cause the function *d* to lose some desirable properties.

9 This feature of the preferential attachment model is what makes it particularly difficult to work with analytically.

10 These results are proven on a model with a slightly different generative procedure; we do not find that they yield a particularly good approximation for our experiments, which are conducted at the quite low *n* = 100.

11 In particular, with these parameters, we observe that the empirical probability of generating a disconnected uncorrelated random graph with these parameters is ~ 0.02%. The preferential attachment section describes in more detail why this exact value is chosen.

12 This is not hard to show; see e.g. [19], Sec III A 1. Furthermore, the *j*^th^ moment of the density gives the expected number of paths of length *j* in the graph.

13 The Python library `NetComp` further simplifies the application of these tools to practical problems; see the appendix for more details.

14 See below regarding the implementation of exact and approximate forms of DeltaCon and the resistance distance.

